# A single computational objective can produce specialization of streams in visual cortex

**DOI:** 10.1101/2023.12.19.572460

**Authors:** Dawn Finzi, Eshed Margalit, Kendrick Kay, Daniel L. K. Yamins, Kalanit Grill-Spector

## Abstract

Human visual cortex is organized into dorsal, lateral, and ventral streams. A long-standing hypothesis is that the functional organization into streams emerged to support distinct visual behaviors. Here, we compare neural network-based computational models against a massive fMRI dataset to investigate why visual streams emerge. We find that a self-supervised Topographic Deep Artificial Neural Network (TDANN), which encourages nearby units to respond similarly, better captures brain responses, as well as spatial segregation and functional differentiation across streams, than DANN models trained for stream-specific visual behaviors. These findings challenge the prevailing view that streams evolved to separately support different behaviors and suggest instead the possibility that functional organization can arise from a single principle: learning generally useful visual representations subject to local spatial constraints.

---

A central question in neuroscience is why the visual system is spatially and functionally organized the way it is. The human visual system is comprised of extensive neural machinery, including more than two dozen visual areas organized into three anatomically distinct hierarchical processing streams, spanning the occipital, temporal, and parietal lobes [1, 2]: (1) the ventral stream, which ascends from early visual cortex (EVC: V1, V2, V3) in the occipital lobe to the inferior aspects of the temporal lobe; (2) the lateral stream, which ascends from EVC through lateral occipito-temporal cortex to the superior temporal sulcus (STS); and (3) the dorsal stream, which projects superiorly from EVC along occipito-parietal cortex. However, it remains unresolved why the visual system is organized into multiple processing streams.

The reasons why the visual system is functionally organized into streams have been studied for decades across empirical [3, 4, 5, 6], information-theoretic [1, 2], and computational [7, 8] approaches. Three key ideas have emerged from these investigations. First, to support complex visual behaviors, the visual system may decompose complex operations into a sequence of simpler computations distributed hierarchically across processing stages [9, 1, 2, 10, 11]. Second, to enable rapid and efficient processing, computations supporting independent visual behaviors, such as recognizing an object (e.g., a car), determining what’s its doing (e.g., approaching), and gauging its distance, are thought to be done in parallel by distinct streams [3, 4, 12, 13, 14]. At the same time, the precise computations supported by each stream remain an active topic of debate, particularly for lateral and dorsal visual streams. The ventral stream is hypothesized to be involved in visual recognition [3, 4] and categorization [15, 16, 17], whereas the lateral stream is hypothesized to be involved in processing visual dynamics, actions, multimodal information (e.g., language), and social information [12, 13, 14, 18], and the dorsal stream is hypothesized to be involved in spatial processing, spatial attention, and visually guided actions [3, 4]. Third, physical constraints, such as fitting the brain into the skull [19] and a need for fast processing, generate a bias for short-range connections [20, 8, 1, 2], which, in turn, may encourage nearby neurons to respond similarly [1, 21, 22, 23]. Indeed, nearby neurons tend to exhibit correlated responses, evident in the many topographic maps across visual cortex, ranging from retinotopic maps [24, 25, 26] in which nearby neurons in cortex process nearby regions of the visual field, to clustering of neurons that selectively respond to specific categories (e.g., faces [27, 28] or words [29]) in high-level visual cortex.

Advances in deep artificial neural networks (DANNs) over the past decade have enabled theories to be transformed into concrete computational models that are trained with images and videos as inputs and tested against real neural data, revolutionizing computational and theoretical neuroscience [30, 17, 31, 32, 33, 34, 35, 36, 37, 38]. To date, the majority of vision-DANN research has focused on understanding the function of the ventral stream, which is empirically the best understood. The main insights from this research are four-fold: (i) feedforward DANNs predict responses across the ventral stream, with earlier layers in DANNs matching responses in early visual cortex (e.g., V1), and later layers matching high-level visual cortex (e.g., ventral temporal cortex, VTC, end of ventral stream) [17, 31, 36], (ii) DANNs trained on behaviorally-relevant tasks, such as categorization for the ventral stream, better predict neural responses than other DANNs [17], (iii) many DANN architectures replicate neural responses in the ventral stream, albeit with numeric differences in performance [35, 39], and (iv) topographic organization within a visual area can emerge through wiring-constrained training [22, 40, 41, 42, 43, 44, 45]. However, most of this work has focused either on ventral cortex or on topography within a single visual area, leaving open whether the same modeling ingredients can explain the broader segregation of cortex into ventral, lateral, and dorsal streams.

Here, we compare two computational accounts of stream organization, instantiated in different model families: optimization of different streams for different candidate visual behaviors (multiple behaviors hypothesis) [3, 46, 47, 48, 49, 17], and learning useful visual representations under spatial constraints to minimize wiring (spatial constraints hypothesis) [19, 8, 1, 2]. Although these model families are not mutually exclusive, comparing the behavior and spatial constraints models allows us to test whether optimization for visual behaviors is necessary for a stream-like organization to emerge. In this framework, behavior-optimized models serve as candidate computational instantiations of prominent theories of stream-specific function, rather than an exhaustive test of all possible behavior-based accounts. To make comparisons across model families possible, we focus on image-computable models. Accordingly, broader behaviors such as navigation, visually guided action, and social interactions are represented through image-computable proxies rather than full closed-loop agentic models. We first establish that the empirical brain data that will be used for model comparison exhibits differentiated responses across streams allowing rigorous model testing, then define the candidate model classes and evaluation framework, and finally test which models best capture the functional and spatial organization into streams.

## Results

### Distinct representational structure and category selectivity across visual streams in the Natural Scenes Dataset (NSD)

Before assessing the models, we tested whether the brain data we will use to test the models – the Natural Scenes Dataset (NSD) [50], which measured high-resolution brain responses to 8000-10,000 of natural images in each of eight individuals during a recognition memory task – shows stream-differentiated responses. Using the 515 NSD images seen three times by all participants, we compute representational similarity matrices for regions of interest (ROIs) spanning early, middle, and high-level visual regions, and compare the representational structure across participants and ROIs. Multidimensional scaling revealed a broad hierarchical progression from early visual cortex to mid-level and then high-level visual ROIs, together with a large-scale separation by stream for high-level regions (Fig. 1a). High-level lateral ROIs were separated from a tight ventral cluster, which was in turn largely distinct from dorsal ROIs, indicating systematic differences in representational structure across the three streams. Mapping responses across these participants’ brains also finds better matches among high-level ROIs within the same stream than across streams (Supplementary Fig. S2).

**Figure 1.**
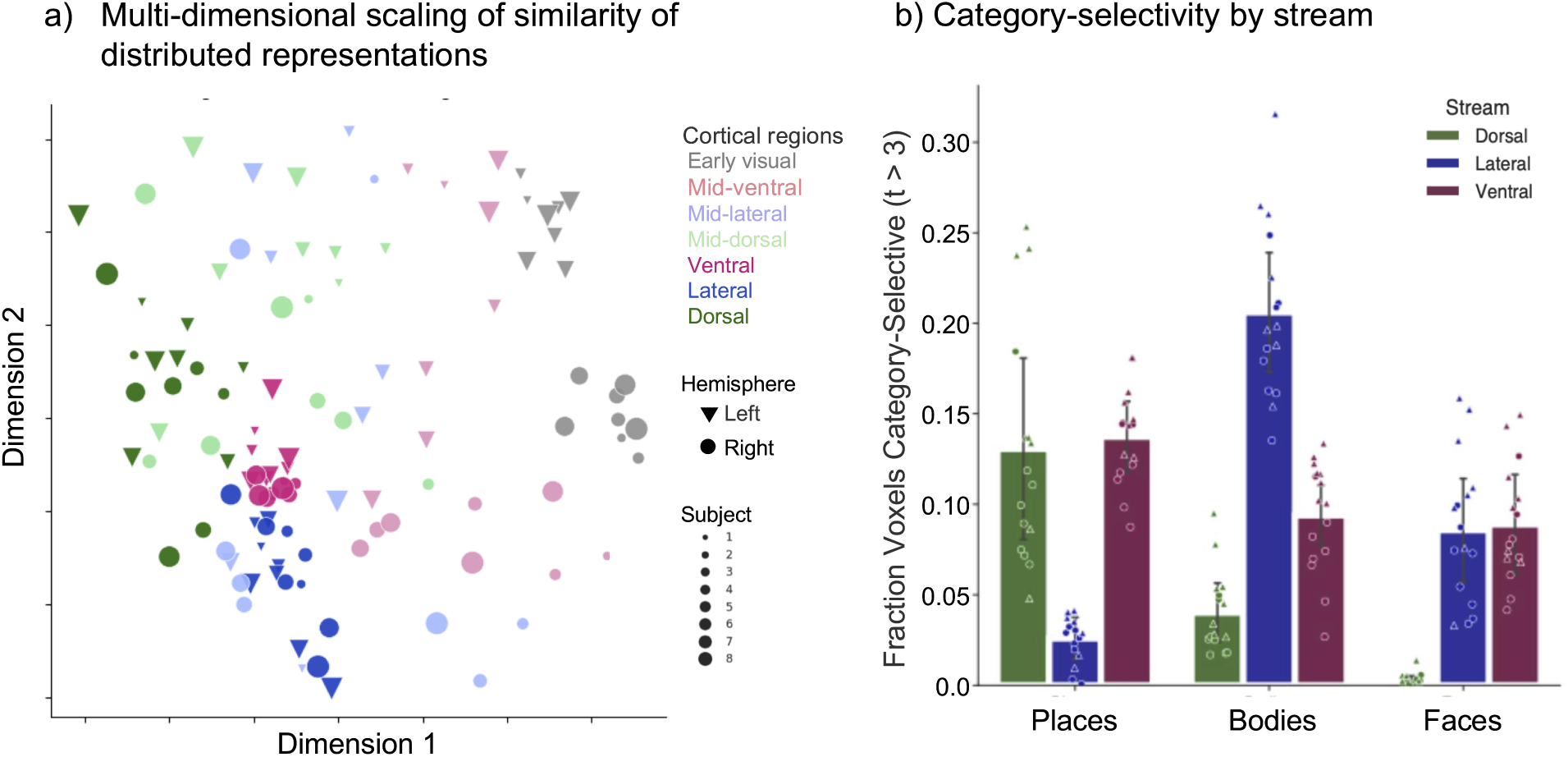
Distinct representational structure and category selectivity across visual streams in the NSD. (a) Multidimensional scaling (MDS) of representational similarity across participants and regions of interest (ROIs) in the Natural Scenes Dataset (NSD). Representational similarity matrices were computed from distributed voxel responses to the 515 images shared across participants. We find a rough hierarchical progression from early visual cortex (EVC) ROIs in the top-right (gray), to mid-level ROIs (light colors, middle), to high-level ROIs in the lower-left. Additionally, there is a large-scale separation by stream for high-level ROIs, rather than by participant or hemisphere, with lateral high-level ROIs (blue) separated and inferior from a tight ventral cluster (red), which is in turn largely distinct from the dorsal ROIs (green, though these show greater between-participant variability). These results indicate that NSD responses exhibit distinct representational structure across dorsal, lateral, and ventral cortex. (b) Fraction of voxels with significant category selectivity (t > 3) for places, bodies, and faces within ROIs consisting of high-level visual cortex in each stream. Category selectivity differs systematically across streams, further demonstrating functional differentiation in the NSD responses, and replicating patterns from the literature. Points represent individual participant–hemisphere combinations. Bars indicate the mean across participants, computed after averaging across hemispheres within each participant. Error bars indicate ±1 SD across participants. Triangles: left hemisphere; circles: right. Green: dorsal; Blue:

Likewise, analysis of brain responses in the same participants to another stimulus set containing controlled images from 10 ecologically-relevant categories (fLoc [51]), finds significant differences in category-selectivity across streams (Fig. 1b) consistent with prior work [16, 52, 53]. Place-selective voxels were more prevalent in ventral (*t*(7) = 21.45, Bonferroni-corrected *p* = 1.1 *→* 10*^→^*^6^) and dorsal (*t*(7) = 6.19, corrected *p* = 4.0 *→* 10*^→^*^3^) cortex than lateral cortex. Body-selective voxels were more prevalent in lateral than ventral (*t*(7) = 13.59, corrected *p* = 2.5 *→* 10*^→^*^5^) and dorsal (*t*(7) = 14.87, corrected *p* = 1.3 *→* 10*^→^*^5^) cortex, and face-selective voxels were more prevalent in ventral (*t*(7) = 8.41, corrected *p* = 5.97 *→* 10*^→^*^4^) and lateral (*t*(7) = 8.93, corrected *p* = 4.03 *→* 10*^→^*^4^) cortex than dorsal cortex. Together, these analyses show that the NSD and fLoc provide functionally differentiated brain responses across streams that will allow us to distinguish models of stream organization.

### Computational framework to instantiate hypotheses for stream organization using deep neural networks

We test which of two model families best predict functional organization into streams: multiple behaviors (MB) or spatial constraints (SC). We instantiate the multiple behaviors hypothesis as the union of three different DANNs, each trained for a stream-hypothesized task. We reason that training different DANNs for different tasks allows optimal task training and encourages learning of maximally different representations suited for each behavior. We use a ResNet-50 backbone [55], as larger models achieve better task performance than shallower models with fewer parameters (e.g., ResNet-18), and previous research has shown a positive correlation between a model’s task performance and its ability to predict neural responses in the ventral stream [17]. In the main multiple behavior model (MB v1), we select tasks that are the computer vision equivalents of behaviors each stream is commonly hypothesized to support - dorsal: object detection [56], lateral: action recognition [57], and ventral: object categorization [55] (Fig. 2a). To test whether results depend on the specific stream functions we model, we implemented two additional variants of stream-specific behaviors. In MB v2, we implemented a vision-language model (CLIP [58]) instead of an action recognition model. This choice was motivated by evidence that the lateral stream extends into posterior superior temporal cortex, which includes multi-modal and speech-/language-related regions [59, 60, 61, 62, 63], recent findings that multi-modal models better capture high-level visual representations than standard supervised networks [64], and theoretical work proposing that the lateral stream may be expanded in humans beyond processing dynamic stimuli to support multi-modal processing such as language [12]. We also changed the detection and categorization models to higher-performing architectures, using DETR [65] as a model for detection and ConvNeXt-T [66] for categorization. In MB v3, we test models for other hypothesized functions of the lateral and dorsal streams, while the ventral stream model remains an object categorization network (ResNet-50), as categorization is the most established candidate function for the ventral stream [17, 39, 31]. For the lateral stream model we implement a human pose-estimation network (DEKR-Pose) [67], motivated by evidence suggesting that the lateral stream is involved in processing body form, motion, and social information [12, 13, 14, 18]. For the dorsal stream model, we implement a monocular depth-estimation network (DepthAnythingV2) [68], motivated by the role of dorsal stream in visually guided behaviors and spatial computations that depend on three-dimensional structure [4, 69]. For all candidate MB models, units from the endpoint layer of the three constituent networks are pooled onto a simulated cortical sheet using the same position-initialization procedure used for topographic models to enable spatial comparison to brain data spanning the union of high-level regions spanning the three visual streams.

**Figure 2.**
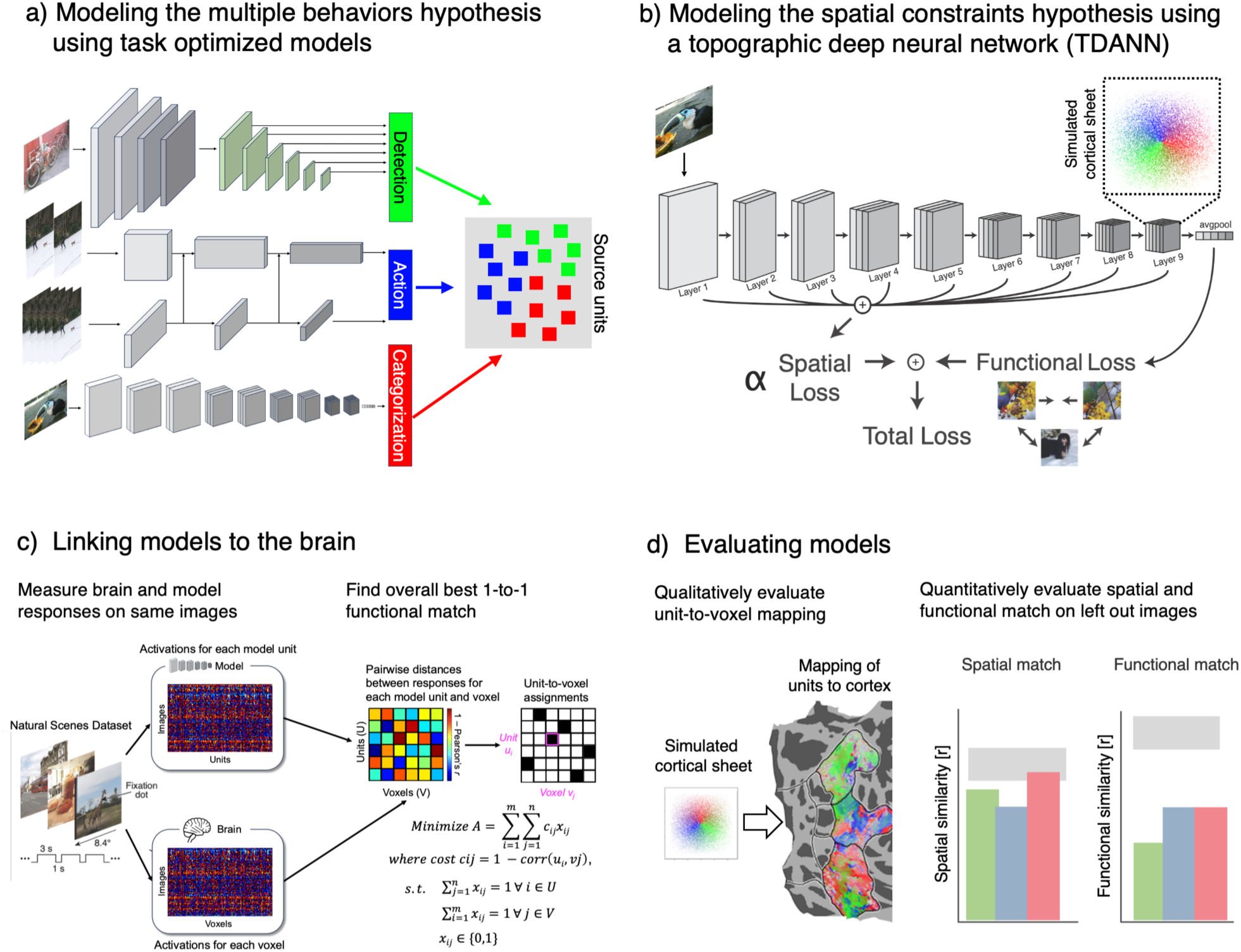
To test competing theories, we use two candidate model classes: Multiple behaviors and Spatial constraints. **(a) Multiple Behaviors (MB):** we implement separate DANNs each trained on a stream-hypothesized task: e.g., detect the object, determine the action, and categorize the object. To compare to the brain, we generate the end region of the MB model by sampling units from the final convolutional layer of each of the three independent DANNs, and pool them into a single combined set of candidate source units on a simulated cortical sheet that will be compared to the union of the endpoint regions of all three streams. **(b) Spatial constraints (SC) models:** we implement Topographic Deep Artificial Neural Networks (TDANNs) [41]. Different than DANNs, during training the TDANN minimizes a composite loss function that has both functional and spatial loss components, with the contribution of the latter controlled by a free parameter *ω*. As for the MB model, to test the spatial constraints model, candidate source units from the final convolutional layer in the TDANN are compared to the union of the endpoint regions of all three streams **(c) 1-to-1 model-to-brain mapping approach.** The goal is to find the overall best functional match between each model unit and each brain voxel. 10,000s of images of natural scenes were presented to each of 8 individuals and candidate models. Responses to identical images are extracted from model units and an individual’s voxels, and correlations between each unit-voxel pair are computed. Correlations are transformed into an initial cost matrix (1 *→* correlation). Using the Hungarian matching algorithm [54] we determine the optimal mapping such that each unit is assigned to a unique voxel and the average cost across all unit-voxel pairings is minimized (black: assignment). **(d) Evaluation.** Models are evaluated by determining spatial correspondence (left) and functional (right) correspondence on 515 left out images seen by all individuals. Spatial and functional correspondences are bench-marked against stream-specific noise ceilings determined by human-to-human mapping of streams using the same approach (light gray).

We instantiate the spatial constraints hypothesis using a topographic DANN (TDANN, [41, 70]), as it implements both a self-supervised learning task (SimCLR [71], Fig. 2b) and spatial constraint during training and has been effective in predicting the functional topography of the ventral stream [41]. In the TDANN, model units in each layer are assigned a position on a 2D simulated cortical sheet, and a spatial constraint is balanced together with contrastive self-supervised learning using separate spatial and task losses during training. The spatial constraint encourages nearby units to have more correlated responses than distant units, and SimCLR encourages two snapshots of the same image (differing in incidental properties such as color or field of view) to have similar representations that are distinct from others [71]. We used SimCLR because it yields broadly useful visual representations [72], an intermediate spatial weight because prior work showed that this regime best reproduces ventral stream topography [41], and a ResNet-18 backbone because it provides a better match between model depth and the number of cortical stages under study [73]. As in prior work, unit positions were initialized using a pre-optimization swapping procedure before joint training.

As controls, we also implement: (i) MB v1 with a ResNet-18 backbone to match the architecture of the spatial constraints model, (ii) another set of spatial constraints models trained on categorization rather than SimCLR, as a large body of research suggests that DANNs trained on categorization better predict responses in the ventral stream [17, 39, 31], (iii) a ResNet-18 DANN trained with SimCLR (no spatial constraint), and (iv) a ResNet-18 DANN trained on categorization (no spatial constraint). These controls allow us to disentangle the effects of backbone architecture, task objective, and spatial loss on each model’s ability to capture stream organization.

### Evaluating functional and spatial correspondence between models and the brain

We use a two-stage approach to evaluate functional and spatial correspondence. First, for each participant, we use the Hungarian algorithm [54] and that participant’s unique image set from the NSD to derive a 1-to-1 mapping between model units in the convolutional end layer and voxels in the union of anatomical ROIs corresponding to the ends of each stream, based on their responses to the same images (Fig. 2c). This 1-to-1 mapping between units and voxels not only provides a window into spatial organization but also provides a more stringent [74, 75] and rigorous [76] testing of DANN models of the brain. Second, using this mapping, we assess spatial and functional correspondence between the candidate model and the brain on an independent set of 515 held-out images seen by all eight NSD participants (5-9% of the total images viewed, depending on the participant; Fig.2d-right). In total, we evaluate 1312 model-to-brain mappings across candidate model types, model seeds, participants, and hemispheres.

We assess spatial correspondence both qualitatively and quantitatively. Qualitatively, we visualize where each model unit maps onto its corresponding voxel on the cortical sheet (Fig. 2d-left). Quantitatively, we measure the similarity between the topography of units on the simulated cortical sheet and that of their assigned voxels in cortex (Methods: Evaluating spatial correspondence; Fig. 2d-right). The spatial similarity metric is the correlation between distances between units on the simulated cortical sheet and distances among their corresponding voxels on the cortical sheet. Spatial similarity is high if nearby units on the simulated cortical sheet are mapped to nearby voxels on the cortical sheet. We assess functional similarity by measuring the correlation between each model unit’s response and that of its corresponding voxel on an independent set of 515 held-out images seen by all participants (Fig. 2d-right).

To assess the maximal performance that models can achieve under this framework, we estimate the noise ceiling of the brain data, hypothesizing that the best model for the human brain is another human brain [77]. We use the Hungarian algorithm and 80% of the 515 shared images for finding the corresponding voxels across participants, and evaluate the spatial and functional correspondence between participants’ brains as above using the remaining 20% of shared images. Most voxels in a given stream of one participant were mapped to the corresponding stream in other participants (Supplementary Fig. S3), validating the mapping procedure and providing stream-specific benchmarks for both spatial and functional correspondence (horizontal gray bars throughout).

### Spatial constraints models provide a better functional and spatial match to the stream organization of visual cortex

We first visualize the topography of unit-to-voxel assignments on the cortical sheet of each participant (Fig. 3a and Fig. S4 for an example participant). Qualitatively, unit-to-voxel assignments for the multiple behaviors (MB) models are noisy, with no clustering by stream (Fig. 3a-left and Fig. S4). In other words, contrary to the prediction of the multiple behaviors hypothesis, units trained on all three tasks are mapped to all three streams and there do not appear to be substantially more units trained on the hypothesized stream behavior mapped to their corresponding stream. For example, categorization-trained units are mapped to all three streams rather than just to the ventral stream as predicted by the multiple behaviors hypothesis. In contrast, the self-supervised spatial constraints model better captures stream topography. Three distinct clusters of units on the simulated cortical sheet map to the three different streams. The model-to-brain mapping is qualitatively more smooth on the cortical sheet than for the multiple behaviors model.

**Figure 3.**
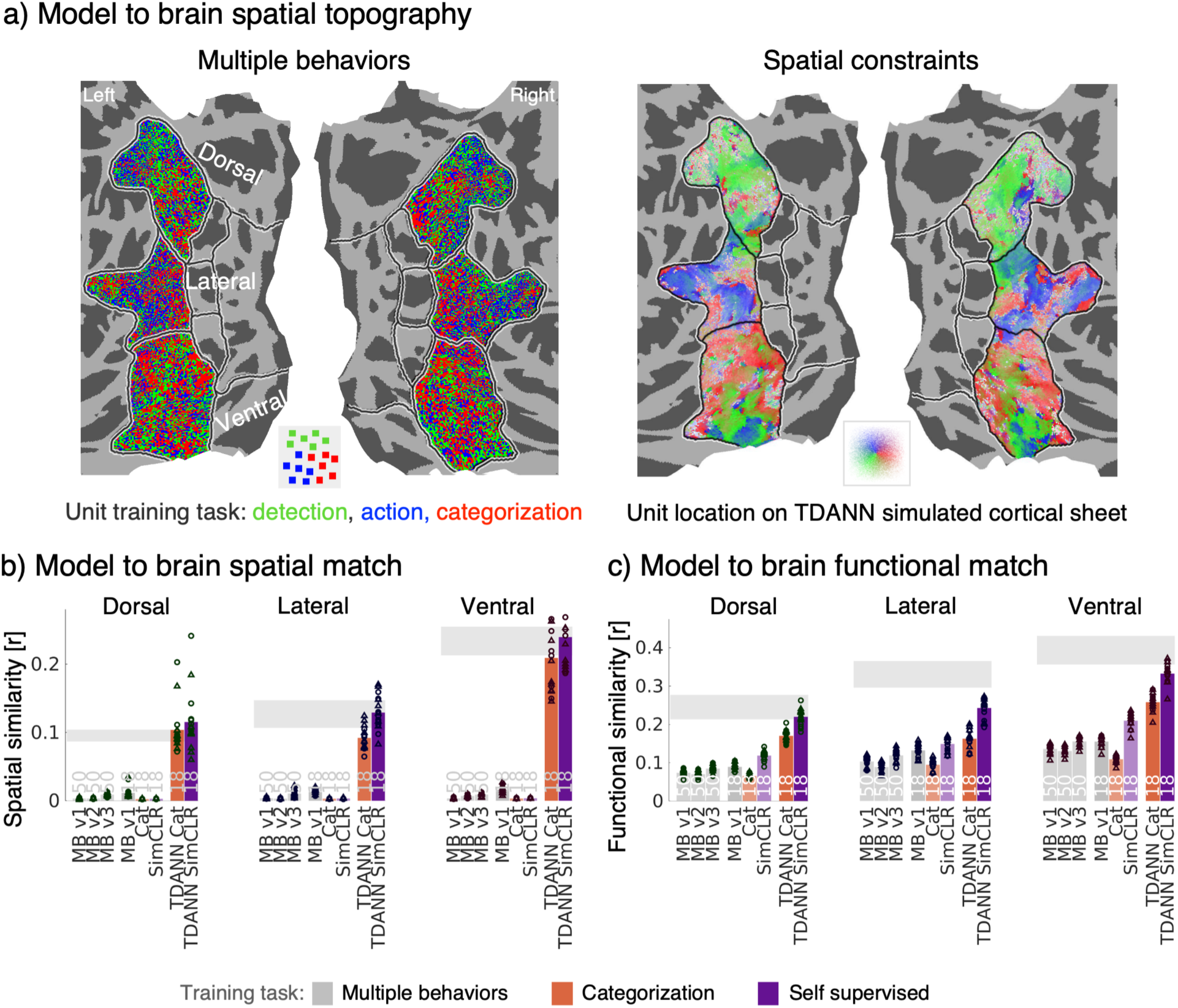
Spatial constraints models better match the spatial and functional organization of the human visual system into streams. **(a)** Unit-to-voxel mapping on an example participant’s flattened cortical surface. *left* : multiple behaviors (MB v1) model, each voxel is colored based on the training task for its assigned unit. *right* : self-supervised spatial constraints model, voxels are colored by location on the simulated cortical sheet of the last convolutional layer. Note that the color scheme for the spatial constraints model represents position and *not* task, as in the left-hand panel. **(b)** Quantification of spatial correspondence. **(c)** Quantification of functional correspondence. Unit-to-voxel correlations on the test set (left-out 515 images). Each participant’s data reflects the average correlation across all voxels for the corresponding stream. For (b) and (c), *colored bars*: mean across participants (N=8) and hemisphere, bars are colored by training task; *numbers*: architecture backbone, 50: ResNet-50, 18: ResNet-18; *horizontal gray bars*: mean noise ceiling across participants and hemispheres ± SD. *Orange*: Supervised (Categorization) and *Purple*: self-supervised (SimCLR) spatial constraints models shown for an optimal weighting of the spatial loss (*ω* = 2.5 for supervised, *ω* = 0.25 for self-supervised). *Acronyms*: MB: multiple behaviors, v1 trained on detection, action recognition, and categorization; v2 trained on more recent models: detection (DETR), language–vision (CLIP), and categorization (ConvNeXt-T); v3 trained with alternative task models for the dorsal and lateral streams: depth estimation (DepthAnythingV2) for dorsal and human pose estimation (DEKR-Pose) for lateral, while retaining the categorization model (ResNet-50) for ventral; TDANN: topographic deep artificial neural network, implementing the spatial constraints hypothesis. Cat: categorization. Source data are provided as a Source Data file.

Quantitative analyses reveal that in all streams, MB models produce close to zero spatial similarity to the spatial arrangement across cortex (Fig. 3b-gray). In contrast, both the self-supervised trained (SimCLR, *ω* = 0.25, Fig. 3b-purple) and categorization-trained (*ω* = 2.5, Fig. 3b-orange) spatial constraints models produce a high spatial similarity to cortex, reaching the human-to-human noise ceiling in all streams (Fig. 3b), with significantly higher spatial correspondence than the MB models (all *p*s *↑* 2.0 *→* 10*^→^*^38^, Supplemental Table 1, Fig. 3b). However, DANNs trained with the same tasks (SimCLR and categorization, respectively) but no spatial constraints produce close to zero spatial similarity between unit positions and their corresponding voxels in visual cortex. This suggests that the spatial constraint during training is necessary for capturing the spatial topography of streams in human visual cortex.

While our results so far suggest that the spatial constraints models capture the stream structure better than the multiple behaviors models, it is possible that the multiple behaviors models still provide a better functional match to the brain than the spatial constraints models. However, even functionally, unit-to-voxel correspondences on the left-out images for the multiple behaviors models are poor and significantly lower than the noise ceiling (all *p*s *↑* 2.5 *→* 10*^→^*^5^, Fig. 3c-gray). For all streams, functional similarity is below *r* = 0.16 for MB v1 (ResNet-50) and MB v3 (ResNet-50), below *r* = 0.15 for MB v2 (ResNet-50), and at or below *r* = 0.17 for MB v1 (ResNet-18). In contrast, the functional correspondence of model units from the self-supervised spatial constraints model to cortex is significantly higher than the multiple behaviors models for all streams (all *p*s *↑* 1.6 *→* 10*^→^*^190^, Fig. 3c-purple, Supplemental Table 2). In fact its functional match is more than double that of the best multiple behaviors models across streams (improvement, dorsal: mean 146% increase, lateral: 84%, ventral: 114%). Notably, the functional correspondence approaches the brain-to-brain noise ceiling (Fig. 3c-gray bars) in both the dorsal and ventral streams. Across all streams, the categorization-trained spatial constraints model (Fig. 3c-orange) also achieves significantly better functional correspondence than the multiple behaviors models (all *p*s *↑* 3.9 *→* 10*^→^*^16^, Supplemental Tables 2) and a better functional match than the task-only categorization trained DANN. However, the functional match of the self-supervised spatial constraints model is significantly higher than the spatial constraints model trained on categorization (pairwise comparisons Bonferroni-corrected, all *p*s *↑* 3.4 *→* 10*^→^*^5^). This difference between the self-supervised and categorization trained spatial constraints models is notable particularly for the ventral stream, as prior research suggests that categorization trained DANNs best predict responses in the primate ventral stream [17, 72]. Thus, across models tested, the self-supervised spatial constraints model provides the best functional and spatial match to the brain (Supplemental Tables 1 and 2). These findings suggest that not only does the spatial constraint in the TDANN change the spatial layout of units, but it also changes their functional properties to be more brain-like.

Because the TDANN explicitly encourages nearby units to respond similarly, one concern is that the smooth spatial clustering observed in Fig. 3a could arise from the architecture rather than from the functional alignment with higher-level cortex. We reasoned that if the spatial smoothness is due to the architecture rather than the functional alignment, then there would be smooth spatial mapping of units from any layer in the TDANN to the different streams. To test this possibility, we repeated the unit-to-voxel mapping using an earlier, V1-like TDANN layer (layer 4 as in [41]). Relative to the optimal late layer, the earlier layer showed substantially weaker spatial correspondence in all three streams: dorsal (0.014 vs. 0.116; *t*(15) = 9.49, *p* = 9.95 *→* 10*^→^*^8^), lateral (0.023 vs. 0.140; *t*(15) = 16.21, *p* = 6.48 *→* 10*^→^*^11^), and ventral (0.017 vs. 0.246; *t*(15) = 20.14, *p* = 2.85 *→* 10*^→^*^12^), as well as weaker functional correspondence (dorsal: 0.092 vs. 0.223; *t*(15) = 30.49, *p* = 6.53 *→* 10*^→^*^15^; lateral: 0.100 vs. 0.252; *t*(15) = 46.81, *p* = 1.13 *→* 10*^→^*^17^); ventral: 0.143 vs. 0.337; paired *t*(15) = 46.91, *p* = 1.09 *→* 10*^→^*^17^; Supplementary Fig. S5). These results show that smooth stream-level organization is not a consequence of the TDANN architecture or the one-to-one mapping procedure, but emerges only when model representations align with higher-level cortical responses.

### Stream-hypothesized task models do not yield stream-specific spatial or functional correspondence

Given the large empirical literature reporting functional differences across streams, we further sought to examine the multiple behaviors models by training task. We considered that our prior analyses, which summarized correspondences across training tasks, might have shown reduced performance due to units being incorrectly mapped to streams. Thus, we evaluated both the spatial and functional correspondence of the multiple behaviors models by individual training task. The multiple behaviors hypothesis predicts that: (i) units trained on a stream’s hypothesized task would be preferentially mapped to the corresponding stream compared to units trained on other tasks, and (ii) units trained on their stream-hypothesized task and assigned to their corresponding stream would demonstrate better functional correspondence than either units trained on other tasks assigned to that stream or units trained on the same task but assigned to incorrect streams.

However, our results largely do not support either prediction. For the dorsal stream, units trained on the stream-hypothesized task of detection map to dorsal voxels at a rate near or slightly higher than chance level (Fig. 4a; mean*±*SD, one-sample t-tests against 33%: SSD = 34.8% *±* 1.5, n.s.; DETR = 35.8% *±* 1.4, *p* = .03; Faster R-CNN = 39.6% *±* 3.0, *p* = 0.008). We observe a similar failure pattern for the lateral stream (SlowFast ResNet-50 = 37.0% *±* 0.8, *p* = 9.2 *→* 10*^→^*^5^; CLIP = 32.5% *±* 1.5, n.s.; SlowFast ResNet-18 = 40.5% *±* 1.5, *p* = 3.3 *→* 10*^→^*^5^). For the ventral stream, units from MB v1 models trained on the stream-hypothesized task of categorization are mapped significantly more than chance to the ventral stream than other streams (ResNet-18: 42.6% *±* 1.7, *p* = 1.5 *→* 10*^→^*^5^; ResNet-50: 41.0% *±* 1.4, *p* = 1.6 *→* 10*^→^*^5^). But for MB v2 model with ConvNeXt-T, surprisingly significantly fewer than chance ConvNeXt-T units are mapped to the ventral stream, despite the categorization training (23.3% *±* 1.4, *p* = 2.2 *→* 10*^→^*^6^). This divergence may reflect ConvNeXt’s attention-inspired design, with larger early receptive fields, LayerNorm-based global context integration, and reduced inductive bias for hierarchical, localized feature encoding, which together may align less closely with the ventral stream’s feedforward and category-selective organization (interestingly, we see a similar decrease in the number of units mapped to the ventral when implementing a ViT-B-32 base architecture control for CLIP, Supplemental Fig. S6). Extending this analysis to MB v3 does not change the conclusion: the depth-estimation model maps significantly below chance level to dorsal cortex (27.7% *±* 2.3%, corrected *p* = 4.5 *→* 10*^→^*^3^), the pose-estimation model does not map selectively to lateral cortex (36.5% *±* 2.7%, corrected *p* = 0.22), and the MB v3 categorization model maps above chance to ventral cortex as in MB v1 (40.3% *±* 1.2%, corrected *p* = 1.4 *→* 10*^→^*^5^). For all MB models, the spatial match remained far below the noise ceiling of 53.8% (*SD* = 1.9%).

**Figure 4.**
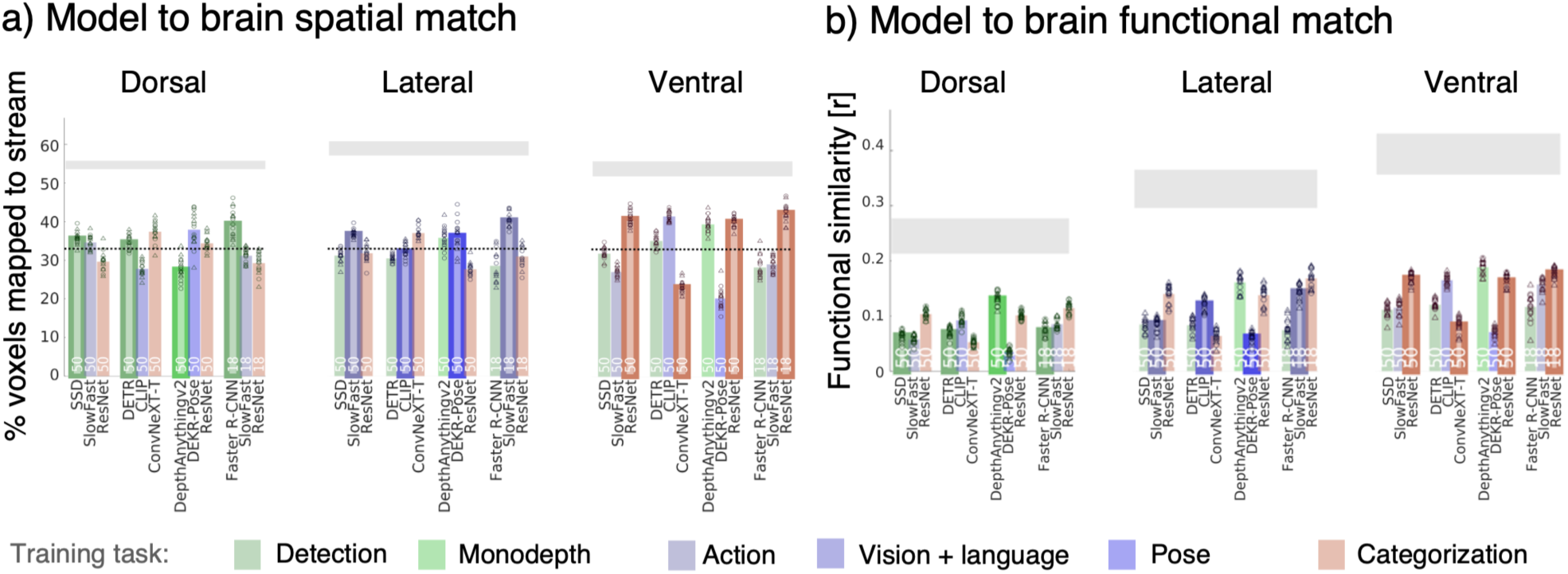
Stream-hypothesized task does not outperform other tasks in predicting brain spatial or functional responses. **(a)** Multiple behaviors model to brain spatial match: proportion of units assigned to stream by training task. Horizontal dashed line: 33 % chance level. **(b)** Multiple behaviors model to brain functional match: functional similarity of units assigned to stream by training task. For (a) and (b) colored bars: mean across participants and hemispheres; bars are colored by training task. light green: monocular depth estimation; blue: action; lavender: vision+language; dark blue: pose estimation; orange: categorization. Outlined bars indicate the stream-hypothesized training. horizontal gray bars: mean noise ceiling across participants and hemispheres ± SD. Source data are provided as a Source Data file.

Functionally, there was likewise no clear dissociation of streams by visual behavior. Across all MB variants and training tasks, units trained on a stream-hypothesized task did not consistently provide the best functional match to that stream. In dorsal, the highest functional similarity among the MB models was observed for the MB v3 depth-estimation model DepthAnythingv2 (mean *±* SD: *r* = 0.131 *±* 0.012), consistent with the proposed role of this model for dorsal processing. In lateral cortex, the best overall MB model remained the categorization-trained ResNet-18 model (*r* = 0.168 *±* 0.016), but among the MB v3 models the depth-estimation model again provided the strongest functional match (*r* = 0.161 *±* 0.016), exceeding both the MB v3 categorization model (*r* = 0.138 *±* 0.018) and the MB v3 pose-estimation model (*r* = 0.062 *±* 0.010). In ventral cortex, the MB v3 depth-estimation model again yielded the highest functional similarity (*r* = 0.189 *±* 0.012), exceeding both the categorization-trained ResNet-18 model (*r* = 0.179 *±* 0.010) and the MB v3 categorization model (*r* = 0.164 *±* 0.012). Thus, while the alternative MB v3 models can modestly improve absolute functional correspondence in some streams, these gains do not produce a clean stream-specific dissociation: depth estimation is the best-performing MB v3 task across all three streams, rather than selectively for dorsal cortex, and overall MB performance remains low relative to the TDANN and the brain-to-brain noise ceiling.

We further examined if the multiple behaviors models failed because the neighborhood-preserving swapping procedure used to initialize unit positions did not converge, or because the task-optimized networks learned highly overlapping representations. To evaluate these possibilities, we tracked both the number of swaps and the neighborhood loss over the course of the swapping optimization and found that both decrease rapidly during early iterations and then stabilize (Supplementary Fig. S7), indicating convergence to a stable solution. We next quantified representational similarity using centered kernel alignment (CKA, [78]) on the end-layer of the three MB v1 ResNet-18 sub-models, since this architecture provided the strongest overall correspondence among the multiple behaviors candidates. We observed moderate similarity between the categorization and action models (CKA = 0.563), but substantially lower similarity between categorization and detection (CKA = 0.114) and between action and detection (CKA = 0.113). Together with the failure of the alternative task choices introduced in MB v2 and MB v3 to recover stream-aligned organization, these analyses suggest that the multiple behaviors framework does not fail simply because of incomplete convergence or a collapsed shared representation among candidate DANNs. While we do not claim to have exhaustively searched the full task space, the present results indicate at minimum that commonly proposed candidate tasks do not readily yield a good match to stream organization under stringent one-to-one unit-to-voxel mapping.

### Two key factors for stream organization: self-supervision and a balanced spatial constraint

We next ask what factors contribute to the emergence of streams in the spatial constraints model. The spatial constraints model has two key components that may affect its performance: the training task and the relative strength of the spatial constraint. Thus, we test spatial constraints models (5 seeds each), varying the training task, contrastive self-supervision (SimCLR) or supervised categorization, and the spatial weighting (*ω*), from *ω* = 0, where the model is essentially a standard ResNet-18 minimizing only the task loss, to *ω* = 25, where the effect of task is dwarfed by the spatial constraint during training. Testing these models against brain data allows us to evaluate: (i) whether there is a broad or narrow range of parameters that gives rise to the stream organization, and (ii) whether similar or different parameters produce spatial and functional correspondences between models and the brain.

We first visualized the spatial topography of spatial constraints model units mapped to their corresponding voxels on the cortical sheet as a function of training task and spatial weight (Fig. 5a). Spatial constraints models trained with no spatial weight (*ω* = 0.0) generate unit-to-voxel mappings which have no spatial structure for both self-supervised and supervised training tasks. I.e., when there is no spatial constraint during training, no functional topography develops in the spatial constraints model’s simulated cortical sheet (Fig. 5a, Supplemental Fig. S8 and S9) and consequently the spatial correspondence is noisy and poor. This suggests that the retinotopic initialization and pre-optimization of units on the simulated cortical sheet is insufficient for the emergence of stream structure.

**Figure 5.**
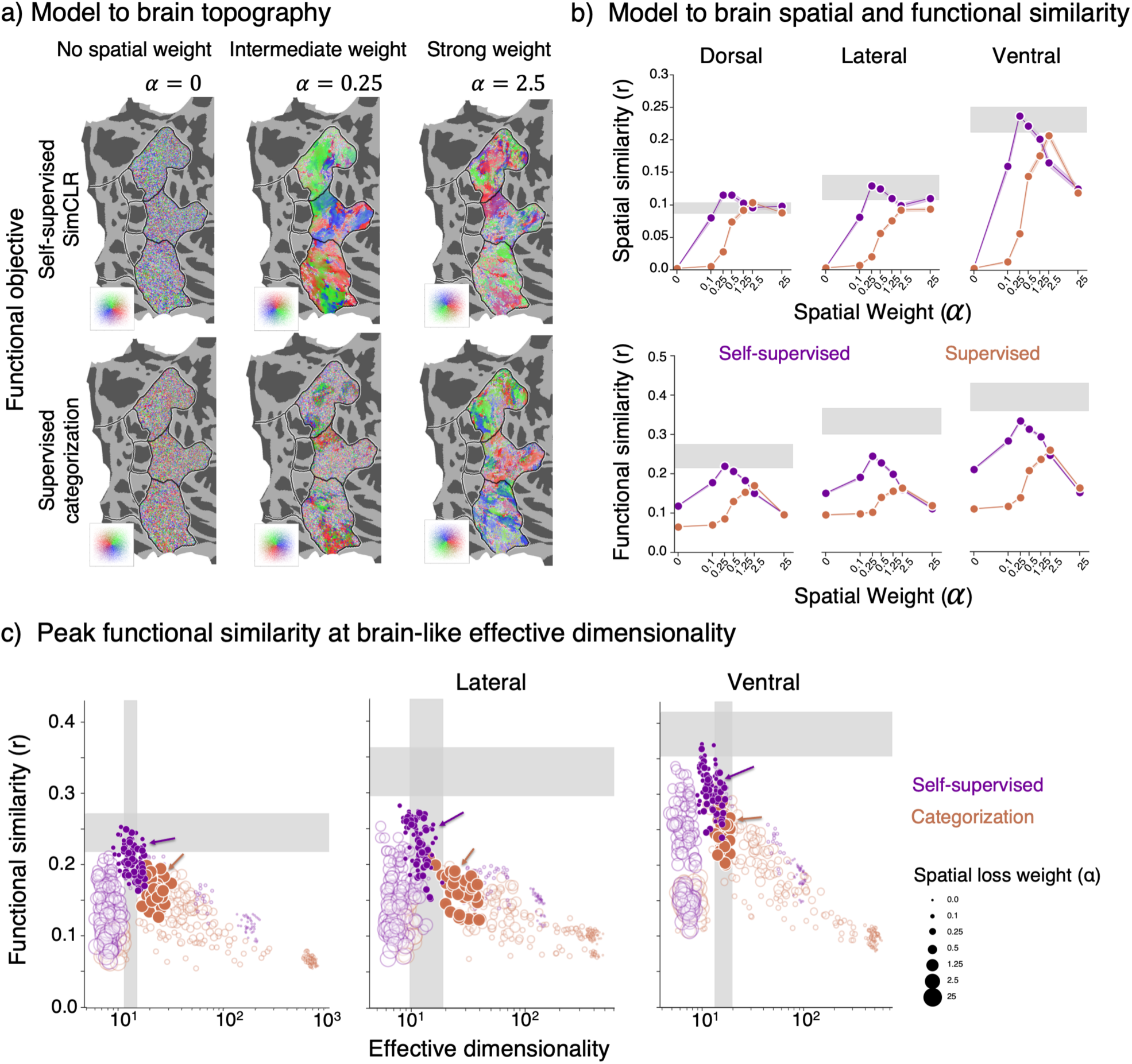
Both self-supervised training and a mid-weight spatial constraint are key to predicting the functional organization into streams. **(a)** Mapping between spatial constraints models and the brain for an example flattened right hemisphere. Voxels are colored by location on the simulated cortical sheet (see inset). Left: spatial constraints models trained with no spatial weighting (*ω* = 0); Middle: spatial constraints models trained with *ω* = 0.25, best functional similarity for self-supervised spatial constraints models; Right: spatial constraints models trained with *ω* = 2.5, best functional similarity for spatial constraints models trained on categorization. **(b)** Across all three streams, the spatial and functional match to brain is best for self-supervised spatial constraints models trained with 0.25 *↑ ω ↑* 0.5. *Top row* : Spatial similarity between unit-voxel pairings per stream. Data averaged over unit-voxel pairing for lowest third of distances. *Bottom row* : average functional similarity (correlation) between unit-voxel pairings per stream. Values are averaged across participants, model seeds and hemispheres; *shaded region*: SD across participants. *Purple*: self-supervised spatial constraints model (SimCLR); *Orange*: supervised spatial constraints model (categorization). *Horizontal gray bar* : noise ceiling, mean ± SD. **(c)** Spatial constraints models with the most brain-like functional similarity also have the most brain-like effective dimensionality. *Horizontal shaded gray bar* : brain-to-brain functional similarity, averaged across participants and hemispheres (*±*SD across target participants). *Vertical shaded gray bar* : effective dimensionality of brain responses by stream, averaged across participants and hemispheres (*±*SD across participants). *Each circle*: one model seed and participant combination, averaged across hemispheres. Circles are colored by training task, and sized by spatial weight (see legend). *Arrows, bolded circles*: models with the highest functional correspondence (b) for each of the spatial constraints training tasks (0.25 *↑ ω ↑* 0.5: self-supervised; *ω* = 2.5: categorization). Source data are provided as a Source Data file.

For spatial constraints models trained with the self-supervised contrastive task, the clearest stream structure is evident when trained with an intermediate spatial weight of *ω* = 0.25, with each contiguous third of the simulated cortical sheet largely mapping to a distinct stream (Fig. 5a-top); this structure is also visible using a continuous spatial gradient (Supplemental Fig. S8, see Supplemental Fig. S9 for corresponding visualizations on the model simulated cortical sheet). But when the spatial weight is increased to *ω* = 2.5 the stream structure is diminished. In contrast, for spatial constraints models trained on a supervised categorization task, training with a stronger spatial weight (*ω* = 2.5) is required for the stream structure to emerge (Fig. 5a-bottom).

Quantitative evaluation of the spatial match to the brain reveals that across all three streams, self-supervised spatial constraints models (purple) provide a better spatial match than supervised (orange) spatial constraints models (Fig. 5b-top, Supplemental Tables 3 to 5). Additionally, spatial correspondence varies significantly with the level of the spatial constraint and there is a significant interaction between the spatial constraint and training task (Supplemental Tables 3 to 5). Across all streams, self-supervised spatial constraints models achieve peak spatial correspondence at 0.25 *↑ ω ↑* 0.5 (Fig. 5b-top), with performance reaching the human-to-human noise ceiling. In contrast, the categorization trained spatial constraints model produces peak spatial correspondence at *ω* = 2.5.

Quantification of the functional similarity between the spatial constraints models and the brain across training tasks and spatial weights mirrors the spatial correspondence, with significantly better functional correspondence for self-supervised than supervised training and a significant task by spatial weight interaction (Fig. 5b-bottom, Supplemental Tables 6 to 8). Across all three streams, spatial constraints models trained with self-supervision and an intermediate spatial weight (0.25 *↑ ω ↑* 0.5) generate the best functional match to the brain. The functional match of the best spatial constraints model approaches the noise ceiling in the dorsal and ventral streams. In contrast, across all streams, spatial constraints models trained on categorization require a larger spatial weight (*ω* = 2.5) to functionally match responses, and even for the ventral stream do not provide a better functional match to the brain than the self-supervised spatial constraints models. Together these analyses reveal that across all three streams, self-supervised spatial constraints models with an intermediate spatial weight (0.25 *↑ ω ↑* 0.5) provide the best spatial and functional match to the brain, suggesting that both the appropriate objective function and spatial constraints during training are needed to generate the functional organization into streams.

What might explain this high functional match between self-supervised spatial constraints models with a balanced spatial weight and the brain? One potential factor lies in the dimensionality of model and brain representations. A characteristic of visual cortex as a computational system is that the dimensionality of its representational space, that is, its effective dimensionality, strikes a balance between efficiency and robustness [79, 80]. Information encoding is most efficient when stimulus responses are uncorrelated and high-dimensional, but most robust when responses are correlated and low-dimensional. The eigenspectra of visual cortex neural responses to natural scenes decay according to a power law with exponent *n ↓* 1 [79, 81], meaning that responses are differentiable and robust to small input perturbations, but sufficiently high-dimensional to encode complex information. Traditional DANNs, on the other hand, exhibit much higher effective dimensionality than neural responses [81, 82], and this higher effective dimensionality is thought to lead to decreased robustness to noise and vulnerability to adversarial attacks [82].

Thus, we next test if spatial constraints models that are more similar to the brain functionally, are also more similar to the brain in their effective dimensionality. We plot the relationship between functional similarity and effective dimensionality of all spatial constraints models tested and compare to the brain. Fig. 5c-gray bars show the functional similarity (horizontal bar) vs. the effective dimensionality of the human brain (vertical bar) in each stream (mean *±* SD: dorsal = 13.2 *±* 1.3; lateral = 14.3 *±* 3.4; ventral = 16.7 *±* 1.9). We find that self-supervised spatial constraints models (Fig. 5c-purple) have lower effective dimensionality than categorization-trained spatial constraints models (Fig. 5c-orange) even as spatial constraints models trained with larger spatial weighting exhibit lower effective dimensionality in general. Comparing the spatial constraints models to the brain data reveals that across all streams the effective dimensionality of the self-supervised spatial constraints models is most similar to the brain at a spatial weight of 0.25 *↑ ω ↑* 0.5 (Fig. 5c-purple arrows). In contrast, the categorization-trained spatial constraints models exceed the effective dimensionality of the lateral and dorsal streams and are only similar in effective dimensionality to the human ventral stream (Fig. 5c-orange arrows). Strikingly, in all streams, the self-supervised spatial constraints models that have the most brain-like effective dimensionality also have the most brain-like functional correspondence. Consistent with this, TDANNs trained with and without the spatial loss are representationally nearly identical at the earliest shared stage (layer 1 CKA = 0.985), but diverge substantially in the deeper layers used for cortical mapping (cross-weight CKA = 0.66 to 0.79, versus pooled within-weight ceilings of 0.87 to 0.95; Supplementary Fig. S10). This pattern suggests that the spatial loss does not simply remap a fixed representation onto the cortical sheet, but progressively alters the higher-level representations most relevant for matching cortical topography and functional responses.

### Observed functional differentiation between streams emerges organically from spatial constraints

Our findings suggest that visual processing streams can emerge in a network that learns via a single, self-supervised task, under a spatial constraint to minimize wiring. Nonetheless, empirical findings reveal multiple functional differences across visual processing streams [3, 47, 14]. Is the emergence of streams from a single training task at odds with functional differentiation across streams?

To gain insights into this question, we test the extent to which spatial constraints model units assigned to different streams may develop stream-relevant functional properties, focusing on three key functional characteristics that have been documented to differ across streams: task specialization, category-selectivity, and receptive field properties.

Firstly, we examine task specialization of units mapped to ventral and dorsal stream voxels, as a large body of work suggests that the ventral stream is involved in visual categorization [17, 15, 16] while the dorsal stream is more important in determining object location [3] even as both position [83, 52] and category [84, 16, 85, 86] can be decoded to some degree from both dorsal and ventral streams. Thus, we test if it is possible to decode from spatial constraints model units assigned to the dorsal and ventral streams, object position [83] and object category (a 1000-way Imagenet categorization task [87, 71]). We hypothesize that if functional differentiation emerges in spatial constraints models, then units assigned to the dorsal stream will outperform ventral units in determining object position and units assigned to the ventral stream [15, 84], will out-perform dorsal stream units on object categorization. We find that this is indeed the case: units mapped to the dorsal stream achieve significantly higher performance than units mapped to the ventral stream on determining object position (mean *±* SD: ventral = 47.8% *±* 0.1%; dorsal = 50.0% *±* 0.4%; *t*(7) = *↔*12.9, *p* = 4.0 *→* 10*^→^*^6^), Fig. 6a). In contrast, units mapped to the ventral stream achieve significantly higher accuracy than units mapped to the dorsal stream on object categorization (mean *±* SD: ventral = 43.7% *±* 0.03%; dorsal = 43.1% *±* 0.2%; *t*(7) = 6.92, *p* = .00023, Fig. 6b). This functional differentiation extends to other tasks and spatial weights (Supplemental Fig. S11).

**Figure 6.**
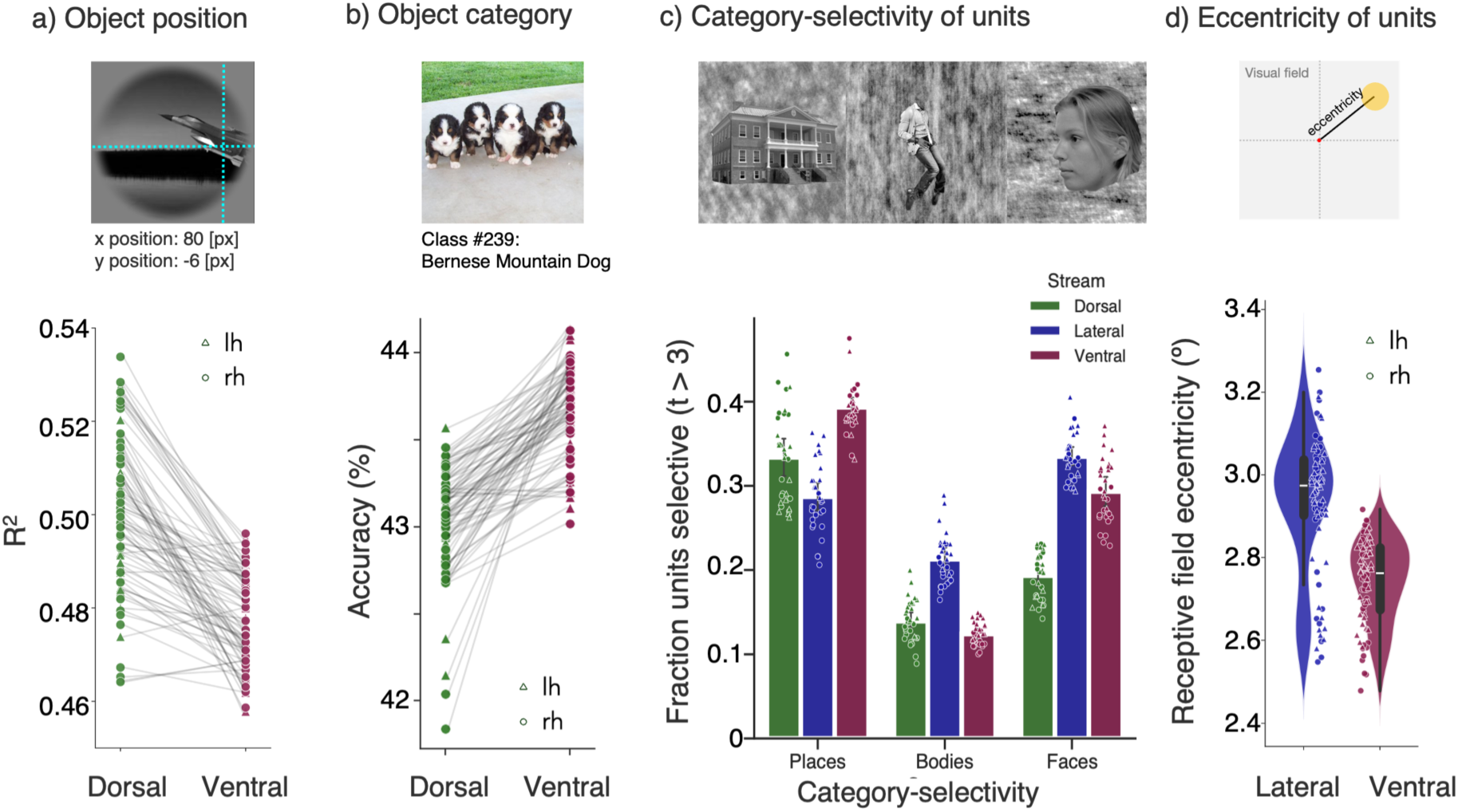
Functional segregation, in alignment with known stream properties, emerges from the spatial constraints model. Data shown is for self-supervised spatial constraints models trained with 0.25 *↑ ω ↑* 0.5. Triangles: left hemisphere; circles: right. Green: dorsal; Blue: lateral; Red: ventral. For (a), (b), and (d), points show individual model seed–participant–hemisphere values, after averaging across spatial-constraint weights (0.25 and 0.5). For (c), points show participant–hemisphere visualization values, after averaging across model seeds and spatial-constraint weights. For all panels, statistical comparisons, where reported, used paired, Bonferroni-corrected, two-sided t-tests across the N=8 participants. **(a)** Top - example of the first transfer task: object position prediction. Bottom - Task transfer performance (*R*^2^) on the object position task (location in pixels of the object’s center [83]) for units assigned to either the dorsal or ventral stream. **(b)** Top - example of the second transfer task: object category prediction. Bottom - Task transfer performance (percentage correct) on the 1000-way object categorization task on the Imagenet validation set for units mapped to the dorsal or ventral stream. **(c)** Top - Example stimuli (images from place, body, and face categories) from the fLoc dataset used to assess category selectivity. Bottom - Relative category selectivity across units assigned to each stream, measured using responses to the fLoc stimulus set. Bars show the fraction of units with significant selectivity (*t>* 3) for each category contrast (places, bodies, faces) within each stream (dorsal, lateral, ventral); mean values across NSD participants, after averaging across model seeds, hemispheres, and the 2 spatial-constraint weights within each participant. Error bars indicate ±1 SD across participants. **(d)** Top - Schematic illustrating the principle of eccentricity. Red dot = center of the visual field. Yellow circle = receptive field. Bottom - Average receptive field eccentricity for face-selective units assigned to the lateral stream or ventral stream. Box plots within violins show the median (center), interquartile range (box bounds: 25th and 75th percentiles), and whiskers extending to the smallest and largest observations within 1.5 times the interquartile range; individual overlaid points show the full data range, including minima, maxima, and any values outside the whiskers. Source data are provided as a Source Data file.

Secondly, as different streams contain different distributions of category-selective voxels (Fig. 1b), we evaluate the category selectivity of stream-assigned units while controlling for low-level image statistics, by measuring units responses on the low-level controlled fLoc stimulus set. Unlike the decoding analyses above, this analysis tests whether the units themselves exhibit distinct selectivity profiles. Indeed, we find that the fraction of units with significant selectivity (*t>* 3) differed systematically across stream assignments (Fig. 6c). Place-selectivity was more prevalent in ventral than lateral assigned units (*t*(7) = 11.17, corrected *p* = 9.2 *→* 10*^→^*^5^); body-selectivity was more prevalent in lateral than ventral (*t*(7) = 14.65, corrected *p* = 1.5 *→* 10*^→^*^5^) and dorsal assigned units (*t*(7) = 8.58, corrected *p* = 5.2 *→* 10*^→^*^4^); and face-selective units were more prevalent in lateral than dorsal assigned units (*t*(7) = 11.35, corrected *p* = 8.3 *→* 10*^→^*^5^) and also more in ventral than dorsal assigned units (*t*(7) = 9.10, corrected *p* = 3.6 *→* 10*^→^*^4^), with no reliable difference between lateral and ventral assigned units (corrected *p* = 0.16). These tuning differences qualitatively mirror the stream-dependent selectivity observed in the brain data (Fig. 1b), indicating that the spatial constraints model captures stream-relevant category tuning in addition to topographic organization and transfer performance.

Thirdly, as population receptive fields (pRFs) in face-selective regions in the lateral stream are more peripheral than those in the ventral stream [52, 88], we test if units mapped to the lateral stream have more peripheral receptive fields than those mapped to the ventral stream. Specifically, we evaluate the mean eccentricity of receptive fields of face-selective units assigned to the lateral and ventral streams based on stimuli spanning 8°x 8°. We find a qualitative model-to-brain correspondence (Fig. 6cd: receptive fields of face-selective units mapped to the lateral stream have significantly higher eccentricity than face-selective units mapped to the ventral stream (mean *±* SD: lateral = 2.93 *±* 0.03; ventral = 2.74 *±* 0.04; *t*(15) = *↔*10.2*,p* = 1.9 *→* 10*^→^*^5^).

Together, these analyses show that even though the spatial constraints model was trained with a single self-supervised task and spatial constraint, units assigned to different streams develop different functional properties, mirroring established patterns in the brain.

## Discussion

We find that a single, biologically plausible computational principle, self-supervised learning of the statistics of visual inputs under a spatial constraint that encourages nearby units to have correlated responses, provides a parsimonious account of both the functional and spatial organization of the human visual system into processing streams. Crucially, both ingredients are necessary: stream structure emerges only when the model combines a self-supervised learning objective with an intermediate spatial weight. Moreover, the spatial constraint that produces the best model-to-brain spatial correspondence also produces the best model-to-brain functional correspondence and most closely matches the effective dimensionality of representations in the human visual system. Thus, a key new idea from our study is that the spatial and functional dimensions of cortical organization are intertwined, and that a local spatial constraint combined with general representation learning can percolate up and generate large-scale structure. Our data also underscore the importance of modeling brain topography alongside function, rather than focusing on function alone, as is done in most current DANN approaches [30, 17, 31, 32, 33, 34, 35, 36, 37, 38], because incorporating spatial constraints yields models that better predict brain function. Taken together, our results suggest that the organization of visual streams may reflect a tradeoff between learning general-purpose representations in a self-supervised manner and conforming to the physical constraints imposed by the size and wiring limitations of the brain.

The success of our spatial constraints model builds upon recent studies of topographic map emergence in the ventral stream [22, 40, 70, 41, 43, 42, 45], which suggest that wiring [22], smoothness [40], and the balancing of spatial and functional constraints [41] can give rise to topographic organization, more energy efficient systems, and can extend to other backbone architectures like visual transformers. Our findings also align with work in mice showing that a dual-pathway model with a self-supervised predictive loss better fits visual cortex data than task-optimized models [89]. At the same time, our results go beyond these studies by showing that explicit optimization of separate pathways for different candidate visual behaviors is not required to reproduce the segregation of visual streams.

We were surprised to find that models based on the multiple behavioral demands hypothesis produced substantially lower functional correspondence with human visual cortex than the spatial constraints model. Based on the cognitive neuroscience literature [3, 4, 90, 12, 13, 14]) our expectation going into the study was that the multiple behavioral demands model would serve as a powerful benchmark model. The performance of the MB models was especially surprising because training tasks that yielded a higher functional match to cortical responses (e.g., categorization) yielded better functional correspondence across all streams, rather than better matching functional responses in their hypothesized cortical stream (e.g., the ventral stream [3, 4, 90, 12, 13, 14]). These findings highlight two important insights. First, they underscore the value of computational modeling as a tool for testing assumptions that may otherwise be taken for granted. Our results show that the common assumption that a model optimized for a behavior associated with a particular cortical stream (e.g., categorization for the ventral stream) should preferentially capture responses in that stream is not necessarily correct. Explicitly implementing and quantitatively testing such theories with DNN models can therefore reveal unexpected outcomes and challenge prevailing intuitions about the functional organization of the brain. Second, while a large body of research has shown that training on categorization yields models that predict ventral stream responses [17, 31, 32, 34, 35, 36], our findings join a more recent body of work demonstrating that training on a variety of other tasks [91, 39], including biologically plausible self-supervised objectives [72, 41], can achieve comparable levels of neural predictivity in the ventral stream.

In contrast, units in the spatial constraints model developed distinct stream-specific functional signatures despite being trained with a single self-supervised objective. This result suggests that functional differentiation can emerge from the interaction of self-supervised learning and spatial constraints, without explicit optimization for separate tasks. Several additional analyses sharpen this interpretation. First, the weaker correspondence of earlier, more V1-like TDANN layers to higher-level visual cortex indicates that stream-like organization does not arise from local smoothness alone. Instead, stream structure emerges in higher layers only once sufficiently rich mid- and high-level visual features have developed. Second, the cross-weight CKA and effective dimensionality analyses indicate that during training the spatial loss does more than impose a spatial arrangement of an otherwise fixed representation on the cortical sheet. Rather, it changes the functional properties of the learned representations, altering their geometry in ways that improve their correspondence with cortical function.

Our conclusions should be interpreted within the scope of the models and data tested here. We have not exhaustively evaluated all possible implementations of the multiple-behaviors hypothesis, nor have we tested models against neural data spanning the full range of human visual behaviors. The NSD data were collected during a single recognition memory task while participants viewed images, thus, our findings should not be viewed as a definitive refutation of all possible multiple-behaviors accounts. Rather, our data provide a proof of existence: explicit optimization for several separate tasks is not necessary for the emergence of stream-like organization. Although the self-supervised spatial constraints model outperforms the multiple-behaviors models, gaps still remain between the predictions of the spatial constraints model and the noise ceiling, particularly in the lateral stream. One possible reason is that both model training and evaluation relied on static images, whereas the lateral stream is thought to be involved in processing dynamic visual information. Thus, although the NSD shows functionally differentiated responses across streams, future work incorporating training on videos, multiple tasks such as next frame prediction, social dynamics, and navigation affordances, and neural data collected during naturalistic video viewing may close the performance gap. Model-to-brain correspondence may also improve by incorporating additional biological features such as long range connections between the ventral and dorsal streams [92, 93, 94] or with downstream areas outside the visual system (e.g., connections between dorsal stream and motor cortex). Computational models also offer a unique opportunity to test causal hypotheses. Future studies could train models on multiple transfer tasks using outputs from all streams, then selectively lesion units assigned to a specific stream, and test whether there are the resulting performance impairments that are consistent with the classic neuropsychological dissociations that motivated parallel-stream theories. Such in-silico lesion experiments would provide a direct test of whether multiple-behaviors models, spatial constraints models, or hybrid accounts best capture the functional organization of the visual system into streams.

Finally, because parallel processing streams exist in other species [90, 89], cortical systems [95, 96, 97, 98], and spatial scales [99, 100], future research can test if the same principles - balancing learning of generally useful representations while encouraging nearby units to have correlated responses – extend to other brain systems. Following the approach established in the present study, this hypothesis could be evaluated by training a topographic deep neural network on diverse sensory and cognitive inputs to test if there are modality-specific or modality-general principles that lead to the emergence of parallel processing streams across the brain.

