## Supplemental Info for "A single computational objective can produce specialization of streams in visual cortex"

### Supplementary Information

### Materials and methods

#### Data availability

The neural data analyzed in this study comes from the Natural Scenes Dataset (NSD) [1] available at http://naturalscenesdataset.org/. Processed data relevant to this study can be found at https://osf.io/qy32x. Additionally, source data are provided with this paper.

#### Code availability

Original code for this study is available at https://github.com/dawnfinzi/spacestream. Code to use and train TDANNs can found at https://github.com/neuroailab/TDANN.

#### Training: Neural network architectures and training tasks

***Multiple behaviors models.*** To test the *multiple behavioral demands hypothesis*, we instantiate model versions of the hypothesis. Each 'multiple behaviors' model is comprised of three sub-models, each trained on a task corresponding to a hypothesized functional goal of a stream. We start by testing the classical view of processing streams using three sub-models, each trained on one of object categorization, action recognition, and object detection (MB v1). We chose these tasks as these are the computer vision equivalents of the proposed behaviors each stream is thought to support (ventral: what is it, lateral: what is it doing, dorsal: where is it). For the object categorization model, we used a ResNet-50 [2] trained on object categorization on ILSVRC-2012 (ImageNet Large-Scale Visual Recognition Challenge [3]). For the action recognition model, we used the SlowFast model architecture [4], which is a dual-pathway network with a 3D ResNet backbone trained on the Kinetics-400 video dataset [5]. Finally, for the object detection model, we used a Faster R-CNN [6] trained on MS-COCO [7]. All three sub-models use ResNet-50 as their base architecture.

Second, we test an alternative (MB v2) using more recent candidate models and replacing action recognition as the hypothesized lateral stream task with fusing language and visual representations, given the lateral stream's proximity to key language areas such as Wernicke's Area and recent work showing Contrastive Language-Image Pretraining (CLIP) models better predict responses in the high-level visual cortex [8]. In this version, for the ventral stream and object categorization, we use the ConvNeXT model [9], again trained on ImageNet, as this next-generation model shows improved categorization performance, as well as being one of the top-performing models on Brain-Score [10]. As mentioned, for the lateral stream, we test the OpenAI trained CLIP [11], where an image encoder and a text encoder are jointly trained to predict the correct pairings of a batch of (image, text) training examples. Finally, for object detection, we use the DETR model [12] trained on MS-COCO. DETR is comprised of a CNN vision backbone and a transformer encoder/decoder architecture. We chose DETR based on the improved task performance [12] and its ubiquitous use in the field. As in the previous version of the multiple behaviors model, we use ResNet-50 or equivalent as a base architecture for all sub-models. For ConvNeXT, this corresponds to the ConvNeXT-T architecture, which is explicitly based on ResNet-50 [9].

Third, we tested an additional multiple behaviors variant (MB v3) to expand the range of candidate functional specializations represented within the multiple behaviors framework. In this version, the ventral stream sub-model remained an object categorization network trained on ImageNet, the lateral stream sub-model was replaced with a human pose estimation network (DEKR, [13]), and the dorsal stream sub-model was replaced with a monocular depth estimation network (Depth Anything V2, [14]). We selected pose estimation as a candidate model of lateral stream function because it emphasizes biological form and body configuration from static images, and is a critical function within social perception [15, 16, 17, 18]. We chose depth estimation as a candidate model of dorsal stream function because it emphasizes three-dimensional scene structure relevant to spatial processing and visually guided behavior [19, 20].

Finally, we implement an additional control using ResNet-18 as the base architecture to more closely match the TDANN architecture, and the number of visual areas in primate cortex [21, 22]. As MB v1 slightly outperformed MB v2 in functional correspondence, this is the version we chose to re-implement using ResNet-18 as a base. For the object categorization and action recognition sub-models, the only change was from ResNet-50 to ResNet-18. However, for the ResNet-18 object detection control model, we used a single-stage object detection network, SSD [23], for greater correspondence with the other multiple behaviors models, and trained on Pascal VOC (2007 and 2012) [24] instead of MS-COCO [7]. While MS-COCO is considered state-of-the-art for object detection and Pascal VOC is an older, smaller dataset, switching to Pascal VOC avoided any potential confound with the images used for the neuroimaging data collection and removed the possibility that the low spatial correspondence for the Resnet-50 based multiple behaviors models was in part due to an artificially inflated match between the object detection model and non-dorsal voxels with high SNR.

We randomly subsampled an equal number of units from layer 4.1 or its equivalent from each network (ignoring any "visually non-responsive" units that did not respond to any of the images) so that the total number of units was equal to the total number of voxels. In order to allow for the most direct comparison against spatial constraints models, the units sampled from the task-trained models were assigned random positions on a two-dimensional simulated cortical sheet. We then followed the same pre-optimization procedure as in Initialization of model unit position: *Stage 2* in order to be able to fairly apply the same mapping algorithm.

***Spatial constraints models.*** We implemented the *spatial constraints hypothesis* using a Topographic Deep Artificial Neural Network (TDANN) architecture. The TDANN is based on the ResNet-18 model architecture, with two key differences: (1) model units are assigned positions on a 2D simulated cortical sheet and (2) the model is trained to jointly minimize a spatial and a task loss. All TDANN models were built using the ResNet-18 [2] base architecture (from the *torchvision* implementation) and trained building on the VISSL framework [25]. ResNet-18 was chosen because it has been shown to achieve strong task performance, accurately predict neuronal responses across the visual system [10], and has roughly comparable number of layers to stages (areas) in the primate visual system [26, 22]. Models were trained for 200 epochs using the ILSVRC-2012 [3] training set, with each model being trained from five different random initial seeds. We optimized the network parameters using stochastic gradient descent with momentum (with $\gamma$ set to 0.9), a batch size of 512, and a learning rate initialized to 0.6, which then decayed according to a cosine learning schedule [27] Models were trained using a self-supervised contrastive objective "SimCLR" [28] or a supervised 1000-way object categorization task.

**Initialization of model unit position** Prior to training with spatial and task losses, model units in each layer were assigned fixed positions in a two-dimensional simulated cortical sheet specific to that layer. The size of the cortical sheet in each layer, and the size of the "cortical zone" used during training (computation of the spatial loss is restricted to units within the same cortical zone), was determined by the presumed correspondence, based on previous work comparing convolutional neural networks (CNNs) and the primate visual system [29, 30, 31], between model layers and human visual areas (see [21] for further details). Positions were then assigned in a two-stage process.

*Stage 1: Retinoptopic initialization* As each layer convolves over the outputs of the previous layer, the resulting responses are structured into spatial grids. To maintain this inherent organization, we assigned each model unit to a specific area of the simulated cortical sheet that aligns with its spatial receptive field.

*Stage 2: Pre-optimization of positions* In CNNs, filter weights are shared between units at different locations, which means that local updates to one unit affect all units with the same filter weights. This global coordination constraint makes it challenging to achieve local smoothness when the units are arbitrarily positioned. To address this, a pre-optimization of unit positions was necessary to identify a set of positions that enables learning smooth cortical maps. We spatially shuffled the units of a pre-trained model on the cortical sheet, so that nearby units had correlated responses to a set of sine grating images. The use of sine gratings is based on studies that show that propagating retinal waves drive development of the visual system in the womb in primates and other mammals [32, 33, 34, 35]

The spatial shuffling works as follows: 1) Randomly select a cortical zone. 2) Compute pairwise response correlations for all units in the zone 3) Select a random pair of units, and swap their locations in the cortical sheet. 4) If swapping positions decreases local correlations (measured as an increase in the Spatial Loss function described below), undo the swap. 5) Repeat steps 3-4 500 times. 6) Repeat steps 1-5 10,000 times.

To verify that the Stage 2 swapping procedure reached a stable solution, we quantified convergence across optimization iterations. For each pre-optimization run, we recorded both the number of accepted swaps and the neighborhood-preserving loss after each iteration block. We then summarized these trajectories across runs and report them in Supplemental Fig. S7. Stabilization of both measures over late iterations was taken as evidence that the swapping procedure had converged rather than terminating prematurely.

**Loss functions** We trained the TDANN models using a weighted sum of two types of loss functions: a task loss, which served to encourage learning of visual representations and a spatial loss, which encourages local correlations in responses to visual inputs. Optimization on the total loss function leads to both successful visual representation learning and minimization of inter-layer wiring length [21].

*Spatial loss* The spatial loss function encourages nearby model units on the simulated cortical sheet to be correlated in their responses to the training stimuli. Specifically, $\mathrm{SL}_l$ is the spatial correlation loss computed for the $l$-th layer and $\mathrm{SL}_l$ is computed on a given batch by randomly sampling a local cortical zone and calculating for pairs of units, (1) correlation (Pearson's $r$) between the response profiles, $(\overrightarrow{R})$, and (2) the stabilized reciprocal Euclidean distances $(\overrightarrow{D})$:

$$\overrightarrow{D} = \frac{1}{(1+\overrightarrow{d})} \quad \textbf{(S1)}$$

where $\overrightarrow{d}$ is the vector of pairwise cortical distances. These two terms are then related as follows:

$$\mathrm{SL}_l = 1 - \mathrm{Corr}\left(\overrightarrow{R}, \overrightarrow{D}\right) \quad \textbf{(S2)}$$

such that $\mathrm{SL}_l$ is minimized when nearby units have correlated responses to the training stimuli.

*Task loss* The task loss ($TL$) is computed from the output of the final model layer. We tested two candidate $TL$s: supervised object categorization cross-entropy loss [36] and the self-supervised SimCLR objective [28]. The SimCLR objective is a contrastive loss function which works by creating two "views", or augmentations, of each image in a batch, using random cropping, horizontal flips, color distortion, and Gaussian blur. These views are passed to the network and the final layer outputs are passed through a 2-layer multi-layer perceptron (MLP), producing a low-dimensional representation of each view which serves as the input to the loss function. The SimCLR loss function then attempts to maximize the similarity of representations for two views of the same source image, while pushing that representation away from all other images in the batch.

**Overview of training** In sum, the model is trained on Imagenet [3] to minimize this total loss, which is the sum of the weighted spatial loss for each layer and the task loss as follows:

$$\text{Total Loss} = \mathrm{TL} + \alpha \sum_{l \in \text{layers}} \mathrm{SL}_l \quad \textbf{(S3)}$$

where $\alpha$ is the weight of the spatial loss component (fixed across all layers), and $\mathrm{SL}_l$ is the spatial correlation loss computed for the $l$-th layer.

The total model training process consists of 6 steps:

1. The model is trained using the task loss only.
2. Positions are initialized to preserve coarse retinotopy in each layer (Stage 1).
3. Positions are pre-optimized in an iterative process that preserves retinotopy while bringing together units with correlated responses to sine gratings (Stage 2).
4. After pre-optimization, positions are permanently frozen.
5. All network weights are randomly re-initialized.
6. The network is trained to minimize the total loss.

#### Testing: Evaluating theories

***Brain data.*** As our neural comparison, we used the Natural Scenes Dataset (NSD) [1], a high-resolution fMRI dataset that densely sampled responses to up to 10,000 natural images in each of eight individuals over the course of 32-40 scan sessions. Full details on data collection and processing can be found in Allen et al. [1]. Briefly, scanning was conducted at 7T using gradient-echo EPI at 1.8-mm isotropic resolution with whole-brain coverage. Images were taken from Microsoft's COCO image database [7], square cropped to 425 pixels x 425 pixels, and presented at fixation at a size of 8.4° x 8.4° for 3 seconds with 1 second gaps in between images.

Data were preprocessed using one temporal resampling (to correct for slice timing differences) and one spatial resampling (to correct for head motion, EPI distortions and gradient non-linearities), resulting in upsampled 1.0mm resolution (temporal resolution 1.0 s). Single-trial beta weights were estimated using a general linear model approach designed to optimize the quality of single trial betas (GLMsingle [37]). We used the single-trial responses from the NSD public data release (beta version b3). We additionally z-scored the betas across images for each voxel and session, averaged across 3 trial repeats, and used cortex-based alignment to align all data to the fsaverage surface. Throughout, the analyzed brain units are fsaverage surface vertices with approximately 1 mm spacing, rather than native 1 mm voxels. We use the fsaverage data in order to standardize the number of vertices in each ROI across participants. Reporting the results, we refer to the brain units of measurement as "voxels" for interpretability, though they are more technically "vertices" given that we are using the fsaverage preparation.

**ROIs** We defined regions of interest (ROIs) for early, intermediate, and high-level visual cortex for each of the three streams based on a combination of anatomical landmarks, noise ceiling estimates (Supplemental Fig. S1), and a constraint to roughly match the number of voxels per stream. We focus only on the high-level visual ROIs for the purposes of this paper and compare the end point of the models to the end points of the processing stream in each brain. However, the full details for drawing all seven (one early, three intermediate and three higher-level) ROIs are included here for completeness and because the high-level ROIs share boundaries with the intermediate ROIs. ROIs were drawn on the fsaverage surface as follows:

Early visual cortex ROI: The early visual cortex ROI was drawn as the union of the V1v, V1d, V2v, V2d, V3v and V3d ROIs from the Wang retinotopic atlas [38]. Additionally, V2v and V2d, as well as V3v and V3d, were connected such that the part of the occipital pole typically containing foveal representations was included in the ROIs.

Intermediate ROIs: Three intermediate ROIs were drawn corresponding to each of the three streams: ventral, lateral and dorsal. All three ROIs border the early visual cortex ROI on the posterior side. The intermediate ventral ROI was drawn to reflect the inferior boundary of hV4 from the Wang atlas [38] and includes the inferior occipital gyrus (IOG), with the anterior border of the ROI drawn based on the anterior edge of the inferior occipital sulcus (IOS). The intermediate lateral ROI was drawn directly superior to the intermediate ventral ROI, with the superior and anterior borders determined as the LO1 and LO2 boundaries from the Wang atlas [38]. The intermediate dorsal ROI was drawn directly superior to that to include V3A and V3B from the Wang atlas.

Higher-level ROIs: Three higher-level ROIs were drawn for each of the ventral, lateral and dorsal streams, bordering their respective intermediate ROIs on their posterior edges. The ventral ROI was drawn to follow the anterior lingual sulcus (ALS), including the anterior lingual gyrus (ALG) on its inferior border and to follow the inferior lip of the inferior temporal sulcus (ITS) on its superior border. The anterior border was drawn based on the midpoint of the occipital temporal sulcus (OTS). The lateral ROI was drawn such that the higher-level ventral ROI was its inferior border and the superior lip of the superior temporal sulcus (STS) was used to mark the anterior/superior boundary. The rest of the superior boundary traced the edge of angular gyrus, up to the tip of the posterior STS (pSTS). The dorsal ROI was drawn to reflect the boundary of the lateral ROI on its inferior edge and to otherwise trace the borders of and include the union of IPS0, IPS1, IPS2, IPS3, IPS4, IPS5 and SPL1 from the Wang retinotopic atlas.

These ROIs, defined by the authors, correspond to the "streams" ROIs included in the NSD public data release. The three higher-level ROI were then trimmed using the prepared noise ceiling maps for beta version b3 and the fsaverage surface [1]. The noise ceiling estimates represent the amount of variance contributed by the signal expressed as a percentage of the total amount of variance in the data, for the average of responses across three trial presentations. An approximate cutoff of 10% was used to guide trimming of the higher-level ROIs, such that we were left with reduced ROIs where all voxels had a noise ceiling $\geq 10\%$ theoretically predictable variance. These ROIs were contiguous and roughly matched in size (number of voxels per ROI right hemisphere: dorsal = 6688, lateral = 6839, ventral = 5638; left hemisphere: dorsal = 6182, lateral = 5849, ventral = 6126).

**Representational structure across visual stream ROIs** To characterize the large-scale organization of visual representations in the target brain data (Fig. 1a), we analyzed responses to the 515 NSD images shared across all eight participants. For each subject, hemisphere, and ROI, we treated the pattern of responses across voxels to each shared image as a distributed response vector and computed the similarity (Pearson's $r$) between the response patterns evoked by all pairs of images, yielding a representational similarity matrix (RSM). We then vectorized the lower triangle of each RSM to obtain one representation vector per subject, hemisphere, and ROI. To compare representational structure across ROIs and participants, we correlated these representation vectors across all subject-hemisphere-ROI combinations, normalizing by trial-to-trial reliability, to generate a second-order similarity matrix. To visualize this structure, we applied multidimensional scaling (MDS) to the second-order matrix and plotted the first two dimensions.

**Category selectivity by stream in human cortex** To benchmark stream-wise category selectivity (Fig. 1b), we used subject-specific fLoc contrast maps in fsaverage space for places, bodies, and faces for the same subjects, again from the NSD public data release. For each subject, hemisphere, stream ROI, and contrast, we extracted the corresponding voxel $t$ values and computed the fraction of voxels exceeding a selectivity threshold of $t > 3$.

***Model data.*** For each of our DANN models, we extracted features in response to the same set of 73,000 total NSD images seen by participants in the scanner. Features were extracted from convolutional layer 4.1 (layer 9) or equivalent. This layer was chosen as past work shows that it has the best functional correspondence to higher-level visual areas, including on the NSD [22, 21, 39]. For the SlowFast network, features were averaged over the temporal dimension. This results in a matrix of features of the form *number of images* x *number of units*, where the number of units is 7 x 7 x 512 = 25,088, i.e. the total number of units in layer 4.1, for the TDANN models, and 19,164 total subsampled units for the multiple behaviors models (6,388 units per model), which matches the maximum number of voxels per hemisphere. Thus, for all models, the number of source units was of the same order of magnitude as the number of target voxels [40, 41].

***1-to-1 mapping between model units and brain voxels.*** To evaluate the model correspondence to the brain we use a 1-to-1 mapping algorithm mapping between model units and brain voxels. We reason that units in a neural network model abstract neural computations and thus may be a good model for the aggregated neural response of a voxel (i.e., in the Goldilocks zone of computational abstraction [42]). To test this reasoning and validate the approach we first developed the algorithm between brain-to-brain, for two reasons. First, in the case of brain-to-brain mapping we are mapping voxels in one brain to voxels in another brain so this tests by definition a mapping between corresponding structures. Second, if the stream organization is not recovered in the 1-to-1 mapping applied between brain-to-brain then we would not be able to test the experimental hypotheses between model and brain. Once satisfied with the mapping algorithm in the brain-to-brain case, we used that algorithm with as few modifications as possible to map model-to-brain.

For each pair of subjects (source subject mapped to target subject), we used the single-trial z-scored betas for each source voxel and each target voxel in response to the $515$ images shared across all subjects (80% used for assignment, 20% used for evaluation) and computed the correlations between each pair of voxels. This correlation matrix was then transformed into a cost matrix (1 – correlation) and assignment was done on the basis of this cost matrix using the Hungarian algorithm [43], a combinatorial optimization algorithm which solves the assignment problem in polynomial time. This 1-to-1 mapping algorithm, which was purely based on functional correlation, performed significantly above chance in assigning voxels to the correct streams (chance = 33%. Mean $\pm$ SE, 1-sample t-test. Left hemisphere: dorsal = $0.519 \pm 0.006$, $t(7) = 31.4, p = 8.6 \times 10^{-9}$); lateral = $0.559 \pm 0.007$, $t(7) = 28.6, p = 1.6 \times 10^{-8}$; ventral = $0.528 \pm 0.006$, $t(7) = 29.5, p = 1.3 \times 10^{-8}$. Right hemisphere: dorsal = $0.575 \pm 0.004$, $t(7) = 63.5, p = 6.3 \times 10^{-11}$); lateral = $0.620 \pm 0.008$, $t(7) = 35.9, p = 3.3 \times 10^{-9}$; ventral = $0.549 \pm 0.008$, $t(7) = 27.0, p = 2.5 \times 10^{-8}$). Consequently, we apply this 1-to-1 mapping in the unit-to-voxel case with only one change. In the unit-to-voxel case, we were no longer restricted to the shared set of $515$ images as we could extract responses from the models to an arbitrary number of images. As such, we leveraged the full set of unique images (up to $9485$ per individual) to link models to individual brains. The $515$ shared images were then reserved for evaluating theories.

***Total models tested.*** We evaluated 5 instances initialized with different random seeds per each of the spatial constraints models across 2 training tasks (SimCLR and categorization) and 7 levels of spatial weightings ($\alpha$), as well as 4 versions of the multiple behaviors models, each version comprised of 3 candidate sub-models. Each model was then mapped to the 2 hemispheres for each of the 8 participants, totaling 1312 model-to-brain mappings tested.

***Evaluating spatial correspondence.*** We evaluated the spatial correspondence in two ways. First, as our main spatial metric to compare the multiple behaviors models and the spatial constraints models directly, we calculated a spatial similarity metric. The spatial similarity metric measures for each unit-to-voxel pairing, the normalized distances on the model cortical sheet (from that unit to other units) and normalized distances on the actual cortical surface (for the assigned voxel to the other units' assigned voxels). For each unit, this metric is calculated across only the closest 33% of units, to simulate stream boundaries, which has the additional benefit of discounting high distances where there are few pairs. This results in two vectors, where each element in the vector is the normalized distance between a pair of units (first vector), or between their assigned voxels (second vector), and the correlation between these two vectors is calculated. This calculation is then repeated for all unit-to-voxel pairings and averaged. The same calculation was performed in the brain-to-brain case to determine the actual cortical level of spatial similarity. We report the average spatial similarity across unit-to-voxel pairings.

For the multiple behaviors models, we additionally evaluated spatial correspondence by calculating the percentage of voxels in each stream that were assigned to the sub-model corresponding to that stream's hypothesized function within a given MB variant. If the hypothesized tasks mapped perfectly to their candidate streams, we would expect to see, for example for MB v1, 100% correspondence between dorsal and the sub-model trained for object detection, 100% correspondence between lateral and action recognition, and 100% correspondence between ventral and object categorization. Chance performance is 33%.

Additionally, for visualization purposes only, in the case of spatial constraints models we divided the TDANN simulated cortical sheet into three sections (candidate stream partition scheme), assuming a log-polar transform, and calculated the highest percentage of voxels that match the partition scheme across candidate partitions (reflection and $5°$ rotations). We could then use that color scheme to visualize back on the brain, for each voxel, the location of that voxel's assigned unit on the simulated cortical sheet.

***Evaluating functional correspondence.*** To evaluate functional correspondence between candidate models and the brain, we report the 1-to-1 correlations calculated on the left-out set of 515 shared images, using the unit-to-voxel assignments determined by the mapping procedure. Each unit-to-voxel correlation was normalized by the individual voxel noise ceiling ($r$) of that assigned voxel (see [1] for information on the calculation of the intra-individual voxel noise ceilings in NSD). 1-to-1 correlations were calculated on an individual subject and hemisphere basis for each of the candidate models. The voxel-to-voxel assignments were used to calculate the overall inter-individual i.e. brain-to-brain noise ceiling (correlations evaluated on test set of 20% of the shared images, averaged across 5 splits for each source and target subject combination).

***Linear regression.*** To compare the 1-to-1 mapping results to the commonly used mapping method of linear regression between model units and the brain [30, 44, 10] (Supplemental Fig. S12), we also calculated model to brain correspondence by regressing model responses from the final convolution layer onto individual voxel responses using ridge regression. As in [10], to decrease computational costs without sacrificing performance, we first projected unit activations into a lower dimensional space using a subsample of the ImageNet validation images and retained the first 1000 PCs. Performance was evaluated on a left-out test set (8/9 train, 1/9 test, shared images excluded, 10 splits) for each subject separately. Test $R^2$ for each voxel is normalized by the individual voxel noise ceiling ($R^2$). To evaluate the upper-bound model performance given the shared variance across subjects, we again calculated a brain-to-brain noise ceiling for each stream by repeating the same ridge regression procedure as for model-to-brain but instead using all other subjects' responses to predict the left-out subject (80/20 train-test split using the set of 515 shared images, 10 splits).

#### Early-layer TDANN control

As an internal control, we repeated the main unit-to-voxel mapping analysis using an earlier, V1-like layer of the best-matching self-supervised TDANN (spatial weight $\alpha = 0.25$). Specifically, we extracted features from the 4th layer, `base_model.layer2.0` (same "V1-like" layer analyzed in [21]) and reran the identical mapping, spatial similarity, and functional correspondence analyses on the higher-level ventral, lateral, and dorsal ROIs. This control tests whether smooth mappings arise trivially from the architecture or instead depend on later representations that better match high-level visual cortex.

#### Effective dimensionality

To calculate effective dimensionality, i.e. the dimensionality of the space of how information is represented by a system (also referred to as the latent dimensionality), we considered the responses of the subset of units assigned to each stream by subject. Using the unit activations (in the case of the models) or z-scored betas (in the case of the human subjects), we computed the eigenspectrum of these responses to the MSCOCO images used in NSD. Following [45] and [46], we computed effective dimensionality from the eigenvalues ($\lambda$) as:

$$ED = \frac{\left(\sum_{i=1}^{N} \lambda_i\right)^2}{\sum_{i=1}^{N} \lambda_i^2} \quad \textbf{(S4)}$$

where $N$ is the number of eigenvectors. Intuitively, if the eigenspectrum decays slowly, that means there are many informative dimensions, and the ED, which in words is simply the squared sum of the eigenvalues over the sum of squares of the eigenvalues, will be high. On the other hand, if the eigenspectrum decays rapidly, meaning that information is largely encoded in only a few dimensions, then the ED will be low.

#### Representational overlap across MB v1 task models

To assess representational overlap within the strongest multiple behaviors baseline, we quantified pairwise overlap across the three MB v1 ResNet-18 task-optimized sub-models using linear centered kernel alignment (CKA, [47]), as in [48]. Specifically, we compared the categorization, action, and detection sub-models rather than stream labels per se. For each pair of task models, we extracted activations to a shared subset of 1,000 NSD images from the same late layers used for voxel mapping (`layer4.1` for categorization, `slow.layer4.1` for action, and `backbone.feature_provider.feature_provider.7.1.conv1` for detection), flattened activations across units, mean-centered responses across stimuli, and computed linear CKA. Higher CKA values indicate greater shared representational structure across tasks. We report pairwise CKA values for categorization–action, categorization–detection, and action–detection, and interpret them alongside the corresponding model-to-brain mapping results.

#### Task transfer

We tested performance of self-supervised spatial constraints model units mapped to the dorsal and ventral stream on new tasks - position prediction and object categorization, respectively - associated with each stream. We refer to this as task transfer performance as the model was not trained on any of these tasks and model weights were frozen. Performance on the transfer task was tested for the self-supervised spatial constraints models which best match the brain ($0.25 \leq \alpha \leq 0.5$ in Fig. 6, full results across spatial weightings, additional tasks and including the lateral stream in Supplementary Fig. S11) and each hemisphere of each subject.

***Position task.*** We evaluated the performance of spatial constraints model units assigned to each stream on predicting the vertical and the horizontal locations in pixels of an object center in an image, using the stimulus set from Hong et al., which has also been used in the evaluation of neural network models of the mouse [50] and primate [44, 51] visual systems. This stimulus set consists of 5760 gray-scale images of 64 distinct objects chosen from one of eight categories (animals, boats, cars, chairs, faces, fruits, planes, tables) placed on randomly chosen, realistic background scene images. Object position, pose and size in this stimulus set varied at different levels from no variation, to medium variation and high variation levels.

For each TDANN model and individual subject, smaller "stream models" were created for each of the three streams by selecting the 5000 units assigned to that stream with the highest correlations. We extracted activations from these units and reduced the dimensionality of the activations to 1000 dimensions using principal components analysis (PCA). We used Ridge regression, with the regularization parameter, $\alpha$, cross-validated from

$$\alpha \in [0.01, 0.1, 1, 10] \tag{S5}$$

to predict the vertical and the horizontal locations in pixels of the object center in the image. We performed five-fold cross-validation on the training split of the no- and medium-variation image subsets, consisting of 3200 images, and computed performance on the test split of the high-variation set consisting of 1280 images. Ten different category-balanced train-test splits were randomly selected. We report $R^2$ on the high-variation test set, averaged across the 10 splits.

***Categorization task.*** To evaluate the performance of TDANN units assigned to each stream on a downstream categorization task, we used the 1000-way ImageNet object categorization task [3]. For each model and individual subject, smaller "stream models" were created for each of the three streams as in the position prediction task. A single linear layer was then trained directly from the outputs of those units. The linear layer was trained for 28 epochs of the ILSVRC-12 ImageNet training set (1,281,167 images) with a batch size of 1,024 and a learning rate which was initialized to 0.04 and decreased by a factor of 10 every eight epochs. We report the top-1 performance on the held-out validation set (50,000 images).

#### Model unit selectivity and receptive field properties

To compare category selectivity across units assigned to different streams, and to relate these proportions to the corresponding voxel proportions in the human data, we used the functional localizer (fLoc) stimulus set [52]. fLoc contains stimuli from five categories, each with two subcategories consisting of 144 images each. The categories are faces (adult and child faces), bodies (headless bodies and limbs), written characters (pseudowords and numbers), places (houses and corridors), and objects (string instruments and cars). These stimuli have been previously used to localize and describe category-selective responses in human higher visual cortex in fMRI studies [52, 53, 54], including in the NSD (see Fig 1).

Selectivity was assessed by computing the $t$-statistic over the set of functional localizer stimuli and defining a threshold above which units were considered selective.

$$t = \frac{\mu_{\text{on}} - \mu_{\text{off}}}{\sqrt{\frac{\sigma^2_{\text{on}}}{N_{\text{on}}} + \frac{\sigma^2_{\text{off}}}{N_{\text{off}}}}}, \tag{S6}$$

where $\mu_{\text{on}}$ and $\mu_{\text{off}}$ are the mean responses to the “on” and “off” categories, respectively, $\sigma^2$ are the associated variances of responses to exemplars from those categories, and $N$ is the number of exemplars being averaged over. As in fMRI experiments, units with $t > 3$ were classified as selective. For each category, the “on” set consisted of the two corresponding fLoc subcategories, and the “off” set comprised all remaining stimuli. We applied this analysis to faces, bodies, and places. For each model seed, subject, hemisphere, and stream, we then computed the fraction of mapped units exceeding the selectivity threshold ($t > 3$) for each category and summarized the resulting selectivity profiles by stream.

Previous studies reported that voxels in face-selective regions in the lateral stream are more peripheral than those in face-selective regions of the ventral stream [53]. To test whether this functional feature was also present in the spatial constraints model, we next identified face-selective units in each stream using the same t-statistic analysis, with the "on" categories defined as adult and child faces and the "off" categories as all non-face categories. For each face-selective unit, eccentricity was then calculated based on the unit's (x,y) position from the center of the $7x7$ filter, converted to degrees of visual angle (by multiplying by $8°$ input stimulus / $7$ grid size). We then divided these units based on which stream they were assigned to and report the mean eccentricity across face-selective units for each model seed × subject × hemisphere combination.

### Supplementary tables

**Table 1. Linear mixed-effects model to test effect of candidate model type on spatial correspondence for each of the three streams.** To test if there were differences between candidate models on spatial correspondence with the brain, we used linear mixed-effects models, with fixed effects for candidate model type (intercept denotes multi-behavior candidate model with ResNet-18 base, i.e. MB v1 ResNet-18) and a random intercept for each subject. Model specification was as follows: spatial correspondence $\sim$ candidate model type + 1 | subject. A separate model was run for each of the three streams. Positive values indicate better spatial correspondence than the MB v1 ResNet-18 (first row) and negative values indicate worse spatial correspondence. For example, the $\beta$ coefficient of $-0.008$ for MB v1 ResNet-50 in dorsal indicates that there is only an average decrease of $0.008$ in spatial correspondence for the MB v1 ResNet-50 model relative to the MB v1 ResNet-18 model, while the $\beta$ coefficient of $0.104$ for the self-supervised TDANN indicates an average increase of $0.104$, again relative to the MB v1 ResNet-18 value of $0.011$. Within each stream, Bonferroni correction was applied to the five model-type coefficients, each representing a contrast against the reference MB v1 ResNet-18 model; the intercept was not included in the correction. $\beta$ coefficients are reported ± SE. Significant predictors ($p < .05$) are shown in bold. MB = multiple behavior. SC = spatial constraints. These statistics are related to Fig 3b.

| | | Coefficients±SE | $z$-value | $p$-value | corrected $p$-value |
|---|---|---|---|---|---|
| Dorsal | Intercept (MB v1 ResNet-18) | 0.011±0.007 | 1.714 | 0.087 | |
| | MB v1 ResNet-50 | -0.008±0.007 | -1.227 | 0.220 | 1.00 |
| | MB v2 ResNet-50 | -0.008±0.007 | -1.228 | 0.219 | 1.00 |
| | MB v3 ResNet-50 | -0.003±0.007 | -0.370 | 0.711 | 1.00 |
| | **SC self-supervised** | **0.104±0.007** | **15.324** | **5.3×10$^{-53}$** | **2.7×10$^{-52}$** |
| | **SC supervised** | **0.092±0.007** | **13.613** | **3.4×10$^{-42}$** | **1.7×10$^{-41}$** |
| Lateral | **Intercept (MB v1 ResNet-18)** | **0.013±0.003** | **3.852** | **1.2×10$^{-4}$** | |
| | MB v1 ResNet-50 | -0.009±0.004 | -2.212 | 0.027 | 0.135 |
| | MB v2 ResNet-50 | -0.010±0.004 | -2.377 | 0.017 | 0.087 |
| | MB v3 ResNet-50 | -0.002±0.004 | -0.525 | 0.600 | 1.00 |
| | **SC self-supervised** | **0.116±0.004** | **28.599** | **7.0×10$^{-180}$** | **3.5×10$^{-179}$** |
| | **SC supervised** | **0.079±0.004** | **19.502** | **1.1×10$^{-84}$** | **5.3×10$^{-84}$** |
| Ventral | **Intercept (MB v1 ResNet-18)** | **0.016±0.008** | **2.130** | **0.033** | |
| | MB v1 ResNet-50 | -0.013±0.009 | -1.351 | 0.177 | 0.883 |
| | MB v2 ResNet-50 | -0.009±0.009 | -0.977 | 0.328 | 1.00 |
| | MB v3 ResNet-50 | -0.007±0.009 | -0.758 | 0.448 | 1.00 |
| | **SC self-supervised** | **0.223±0.009** | **23.975** | **5.1×10$^{-127}$** | **2.6×10$^{-126}$** |
| | **SC supervised** | **0.193±0.009** | **20.716** | **2.5×10$^{-95}$** | **1.2×10$^{-94}$** |

**Table 2. Linear mixed-effects model to test effect of candidate model type on functional correspondence for each of the three streams.** To test if there were differences between candidate models on functional correspondence with the brain, we used linear mixed-effects models, with fixed effects for candidate model type (intercept denotes multi-behavior candidate model with ResNet-18 base, i.e. MB v1 ResNet-18) and a random intercept for each subject. Model specification was as follows: functional correspondence $\sim$ candidate model type + 1 | subject. A separate model was run for each of the three streams. Positive values indicate better functional correspondence than the MB v1 ResNet-18 (first row) and negative values indicate worse functional correspondence. For example, the $\beta$ coefficient of $-0.019$ for MB v1 ResNet-50 in ventral indicates that there is an average decrease in the correlation with brain responses of $0.019$ for the MB v1 ResNet-50 model relative to the MB v1 ResNet-18 model, while the $\beta$ coefficient of $0.178$ for the self-supervised SC model indicates a massive average increase in correlation of $0.178$, again relative to the MB v1 ResNet-18's value of $0.155$, meaning that the self-supervised SC model had an average correlation across subjects of $0.333$ to ventral brain responses. Within each stream, Bonferroni correction was applied to the five model-type coefficients, each representing a contrast against the reference MB v1 ResNet-18 model; the intercept was not included in the correction. $\beta$ coefficients are reported ± SE. Significant predictors ($p < .05$) are shown in bold. MB = multiple behavior. SC = spatial constraints. These statistics are related to Fig 3c.

| | | Coefficients$\pm$SE | $z$-value | $p$-value | corrected $p$-value |
|---|---|---|---|---|---|
| Dorsal | **Intercept (MB v1 ResNet-18)** | **0.089$\pm$0.004** | **23.640** | **1.5$\times$10$^{-123}$** | |
| | **MB v1 ResNet-50** | **-0.016$\pm$0.004** | **-4.135** | **3.6$\times$10$^{-5}$** | **1.8$\times$10$^{-4}$** |
| | **MB v2 ResNet-50** | **-0.019$\pm$0.004** | **-5.157** | **2.5$\times$10$^{-7}$** | **1.3$\times$10$^{-6}$** |
| | MB v3 ResNet-50 | -0.005$\pm$0.004 | -1.275 | 0.202 | 1.00 |
| | **SC self-supervised** | **0.130$\pm$0.004** | **34.763** | **9.0$\times$10$^{-265}$** | **4.5$\times$10$^{-264}$** |
| | **SC supervised** | **0.081$\pm$0.004** | **21.552** | **5.1$\times$10$^{-103}$** | **2.6$\times$10$^{-102}$** |
| Lateral | **Intercept (MB v1 ResNet-18)** | **0.132$\pm$0.007** | **20.062** | **1.6$\times$10$^{-89}$** | |
| | **MB v1 ResNet-50** | **-0.029$\pm$0.004** | **-8.135** | **4.1$\times$10$^{-16}$** | **2.1$\times$10$^{-15}$** |
| | **MB v2 ResNet-50** | **-0.043$\pm$0.004** | **-12.149** | **5.8$\times$10$^{-34}$** | **2.9$\times$10$^{-33}$** |
| | **MB v3 ResNet-50** | **-0.013$\pm$0.004** | **-3.646** | **2.7$\times$10$^{-4}$** | **1.3$\times$10$^{-3}$** |
| | **SC self-supervised** | **0.111$\pm$0.004** | **31.053** | **1.0$\times$10$^{-211}$** | **5.2$\times$10$^{-211}$** |
| | **SC supervised** | **0.031$\pm$0.004** | **8.586** | **9.0$\times$10$^{-18}$** | **4.5$\times$10$^{-17}$** |
| Ventral | **Intercept (MB v1 ResNet-18)** | **0.155$\pm$0.006** | **27.244** | **1.9$\times$10$^{-163}$** | |
| | **MB v1 ResNet-50** | **-0.019$\pm$0.004** | **-4.988** | **6.1$\times$10$^{-7}$** | **3.0$\times$10$^{-6}$** |
| | **MB v2 ResNet-50** | **-0.024$\pm$0.004** | **-6.432** | **1.3$\times$10$^{-10}$** | **6.3$\times$10$^{-10}$** |
| | MB v3 ResNet-50 | 0.000$\pm$0.004 | 0.115 | 0.909 | 1.00 |
| | **SC self-supervised** | **0.178$\pm$0.004** | **46.815** | **2.1$\times$10$^{-478}$** | **1.0$\times$10$^{-477}$** |
| | **SC supervised** | **0.103$\pm$0.004** | **27.145** | **2.9$\times$10$^{-162}$** | **1.5$\times$10$^{-161}$** |

To test for the effects of TDANN spatial weightings ($\alpha \in [0.0, 0.1, 0.25, 0.5, 1.25, 2.5, 25]$) and training task (self-supervised simCLR vs. supervised object categorization) on (1) model-to-brain spatial similarity (Fig. **??**b-top) and (2) model-to-brain functional similarity (Fig. 5b-bottom), we ran repeated-measures ANOVAs separately for each stream with the factors spatial weighting and training task. Results are reported in Tables 3-8. Num DF indicates numerator degrees of freedom and Den DF indicates denominator degrees of freedom.

**Table 3. Spatial similarity ($r$) for Dorsal.** Two-factor repeated-measures ANOVA (n = 8; within-participant factors: spatial weight and training task); P values are from upper-tailed F tests with no multiple-comparison adjustment.

| | F | Num DF | Den DF | p-value |
|---|---|---|---|---|
| Spatial weighting | 79.98 | 6.0 | 42.0 | $2.3\times10^{-21}$ |
| Training task | 53.87 | 1.0 | 7.0 | $1.6\times10^{-4}$ |
| Spatial weighting:Training task | 47.56 | 6.0 | 42.0 | $3.7\times10^{-17}$ |

**Table 4. Spatial similarity ($r$) for Lateral.** Two-factor repeated-measures ANOVA (n = 8; within-participant factors: spatial weight and training task); P values are from upper-tailed F tests with no multiple-comparison adjustment.

| | F | Num DF | Den DF | p-value |
|---|---|---|---|---|
| Spatial weighting | 157.95 | 6.0 | 42.0 | $3.6\times10^{-27}$ |
| Training task | 150.65 | 1.0 | 7.0 | $5.5\times10^{-6}$ |
| Spatial weighting:Training task | 78.58 | 6.0 | 42.0 | $3.2\times10^{-21}$ |

**Table 5. Spatial similarity ($r$) for Ventral.** Two-factor repeated-measures ANOVA (n = 8; within-participant factors: spatial weight and training task); P values are from upper-tailed F tests with no multiple-comparison adjustment.

| | F | Num DF | Den DF | p-value |
|---|---|---|---|---|
| Spatial weighting | 162.07 | 6.0 | 42.0 | $2.1\times10^{-27}$ |
| Training task | 242.89 | 1.0 | 7.0 | $1.1\times10^{-6}$ |
| Spatial weighting:Training task | 217.25 | 6.0 | 42.0 | $5.7\times10^{-30}$ |

**Table 6. Functional similarity ($r$) for Dorsal.** Two-factor repeated-measures ANOVA (n = 8; within-participant factors: spatial weight and training task); P values are from upper-tailed F tests with no multiple-comparison adjustment.

| | F | Num DF | Den DF | p-value |
|---|---|---|---|---|
| Spatial weighting | 247.50 | 6.0 | 42.0 | $4.1\times10^{-31}$ |
| Training task | 534.95 | 1.0 | 7.0 | $7.2\times10^{-8}$ |
| Spatial weighting:Training task | 411.76 | 6.0 | 42.0 | $1.2\times10^{-35}$ |

**Table 7. Functional similarity ($r$) for Lateral.** Two-factor repeated-measures ANOVA (n = 8; within-participant factors: spatial weight and training task); P values are from upper-tailed F tests with no multiple-comparison adjustment.

| | F | Num DF | Den DF | p-value |
|---|---|---|---|---|
| Spatial weighting | 221.47 | 6.0 | 42.0 | $3.9\times10^{-30}$ |
| Training task | 628.36 | 1.0 | 7.0 | $4.1\times10^{-8}$ |
| Spatial weighting:Training task | 520.25 | 6.0 | 42.0 | $9.5\times10^{-38}$ |

**Table 8. Functional similarity ($r$) for Ventral.** Two-factor repeated-measures ANOVA (n = 8; within-participant factors: spatial weight and training task); P values are from upper-tailed F tests with no multiple-comparison adjustment.

| | F | Num DF | Den DF | p-value |
|---|---|---|---|---|
| Spatial weighting | 511.53 | 6.0 | 42.0 | $1.3\times10^{-37}$ |
| Training task | 974.32 | 1.0 | 7.0 | $9.0\times10^{-9}$ |
| Spatial weighting:Training task | 768.23 | 6.0 | 42.0 | $2.9\times10^{-41}$ |

#### Supplementary figures

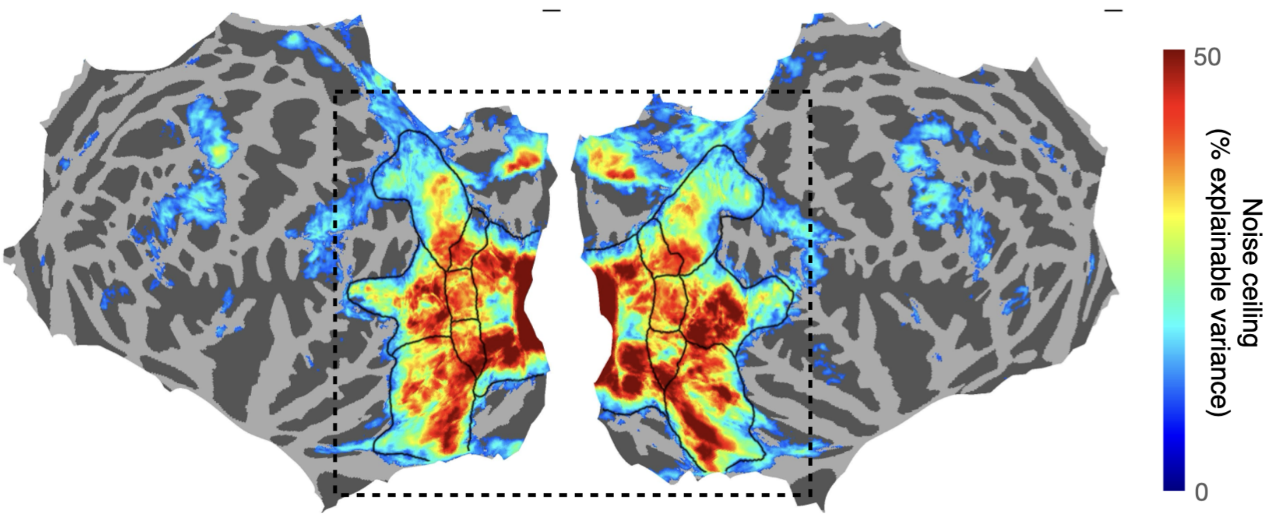


**Figure S1. Voxel-wise noise ceiling estimates and ROI boundaries.** Noise ceiling estimates (% explainable variance) across all image repeats per subject and then averaged across subjects, visualized on the fsaverage surface. Values are thresholded at 10% explainable variance, the cutoff used to guide drawing of the higher-level ROI boundaries. Figure illustrates that, (1) by design, much of the reliable signal is included in the ROIs drawn, and (2) the noise ceiling is high (minimum 10% explainable variance and numerous voxels above 50% explainable variance) across ventral, lateral, and dorsal. Dashed black line: region shown in main text figures.

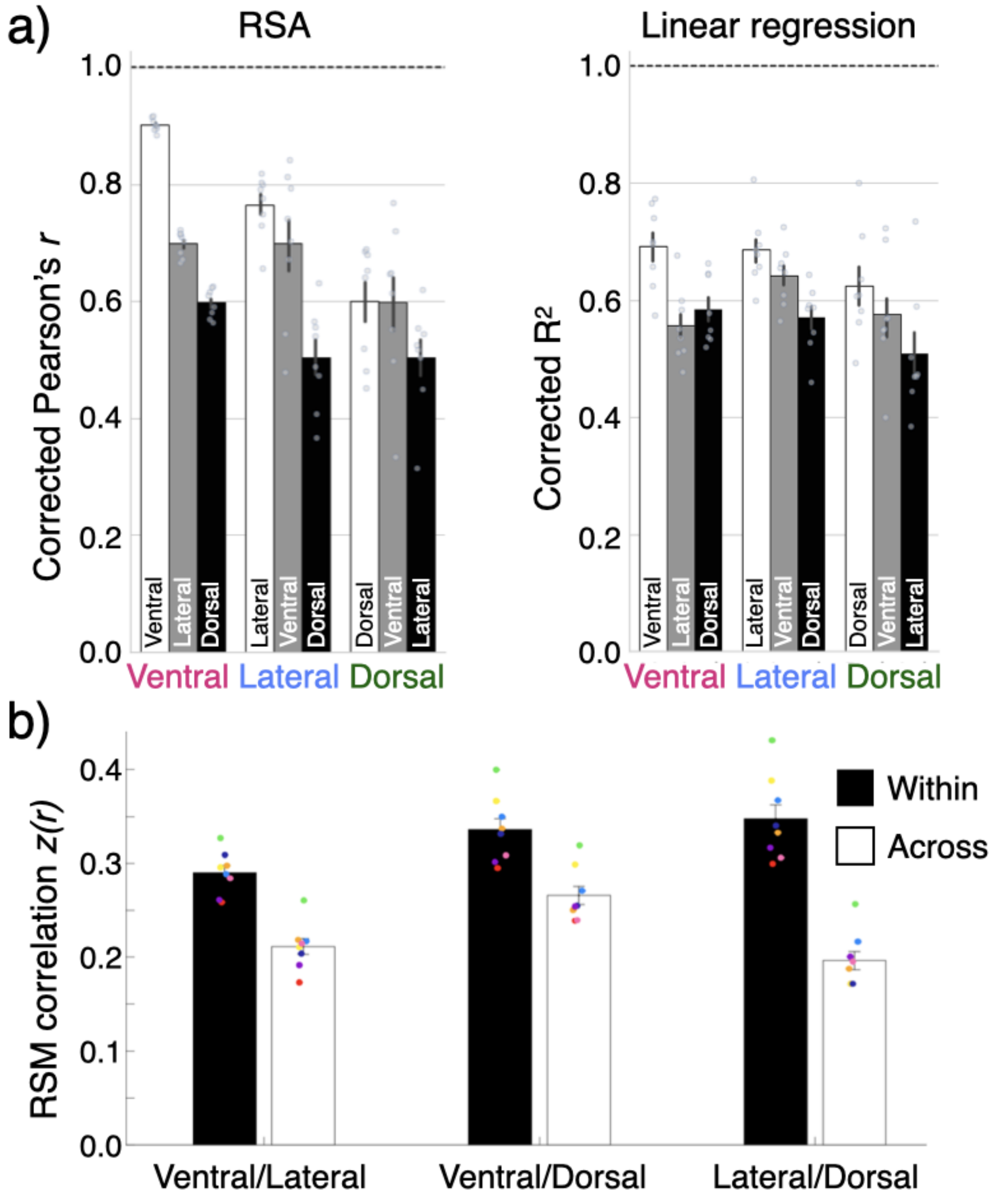


**Figure S2. Additional evidence for representations differing across streams in NSD.** (a) Comparison of ROIs as models of each other, using representational similarity analysis (RSA) and linear regression (Ridge regression). Pearson's $r$ and $R^2$ values are normalized by the respective noise ceilings (NC). Each dot represents a subject. White: within-ROI (i.e. subject-to-subject noise ceiling); Gray and Black bars: ROI X's prediction of ROI Y's responses. (b) To further test whether each stream showed a distinct representational structure, we parcellated cortex into 1000 equally spaced ROIs and then calculated the correlation between each pair of parcels. Each comparison was grouped based on whether both parcels were located within the same stream (black) or whether they were located in two different streams (white), revealing significantly higher correlations within than across streams for this three-stream organization (main effect of within vs. across: $p$=4.19x$10^{-7}$). The difference in parcel correlations within vs. across streams did not simply reflect anatomical proximity, as the neighboring lateral and dorsal streams showed the greatest differentiation. Source data are provided as a Source Data file.

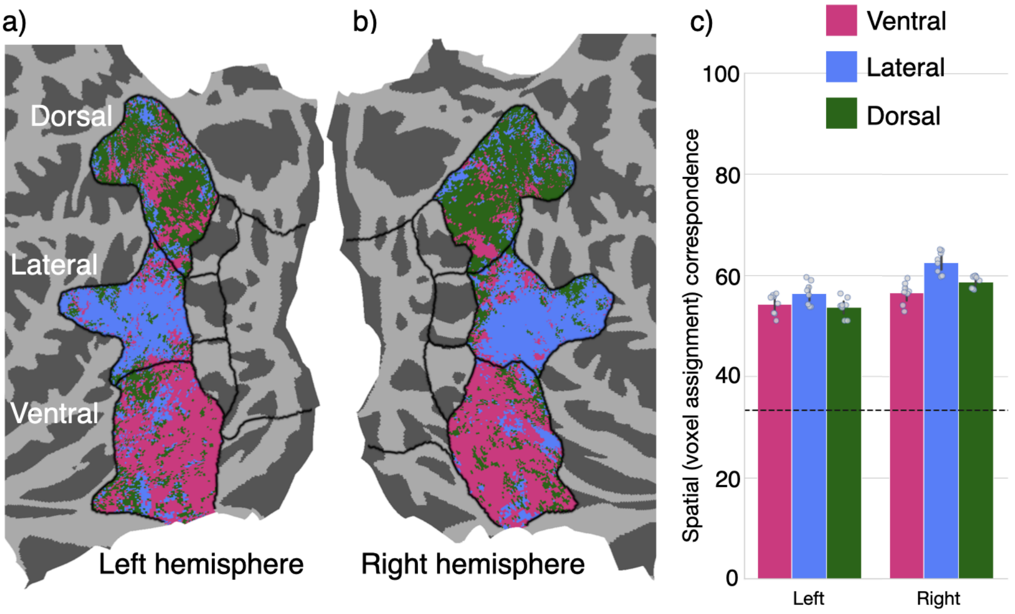


**Figure S3. Validating the 1-to-1 mapping algorithm by testing how well it maps one brain onto another brain.** (a) Left hemisphere voxels in Target (Subj. 2) brain colored by their assignment to streams in Source (Subj. 1) left hemisphere voxels using the 1-to-1 mapping algorithm [43] that matches each unit to a voxel based on functional similarity. (b) Same for the right hemisphere. (c) Quantifying the spatial correspondence between brains achieved using the 1-to-1 mapping algorithm algorithm. Data plotted for high-level ROIs across three streams, averaged across source subjects for each target subject. Color represents stream (pink: ventral, blue: lateral, and green: dorsal). Dotted line: chance level (33%). Error bar: 95% CI, each dot is a subject. Data show that using this purely functional 1-to-1 matching procedure, nearly two-thirds of units are correctly mapped to the same stream between source and target subjects, significantly above chance (all $ps \leq 2.5 \times 10^{-8}$). Source data are provided as a Source Data file.

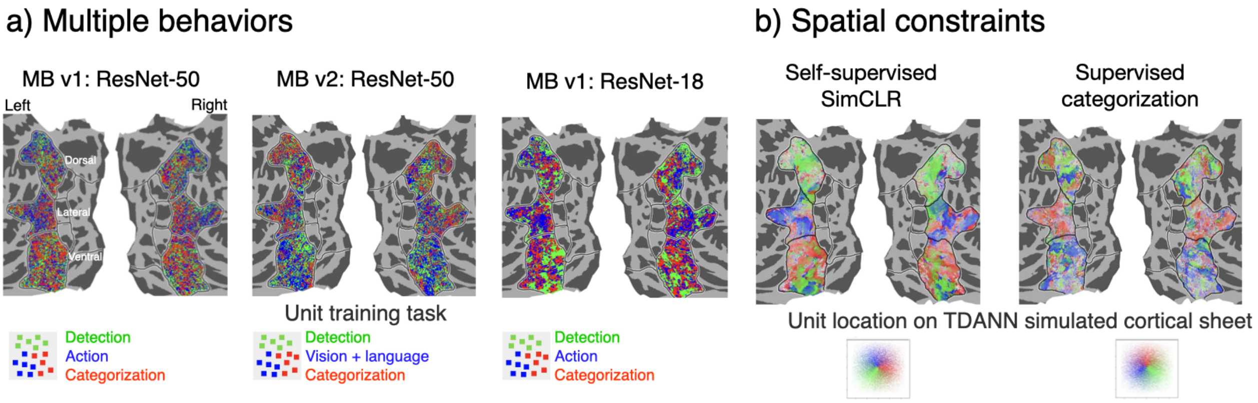


**Figure S4. Mapping between candidate multiple behaviors and spatial constraints models and the brain for an example participant's flattened cortical surface.** Unit-to-voxel mapping visualized on an example participant's flattened cortical surface. (a) multiple behaviors (MB) models, each voxel is colored based on the training task for its assigned unit. (b) spatial constraints (SC) models, voxels are colored by location on the simulated cortical sheet of the last convolutional layer; polar angle (red -> green -> blue) and eccentricity (opacity, which decreases away from center). For each row, the control models are on the right (ResNet-18 base architecture for the multiple behavior models and supervised categorization training for the spatial constraints models). The visualizations for MB v1 ResNet-50 and the self-supervised SimCLR SC model are the same as Figure 3a, replotted here to facilitate comparison.

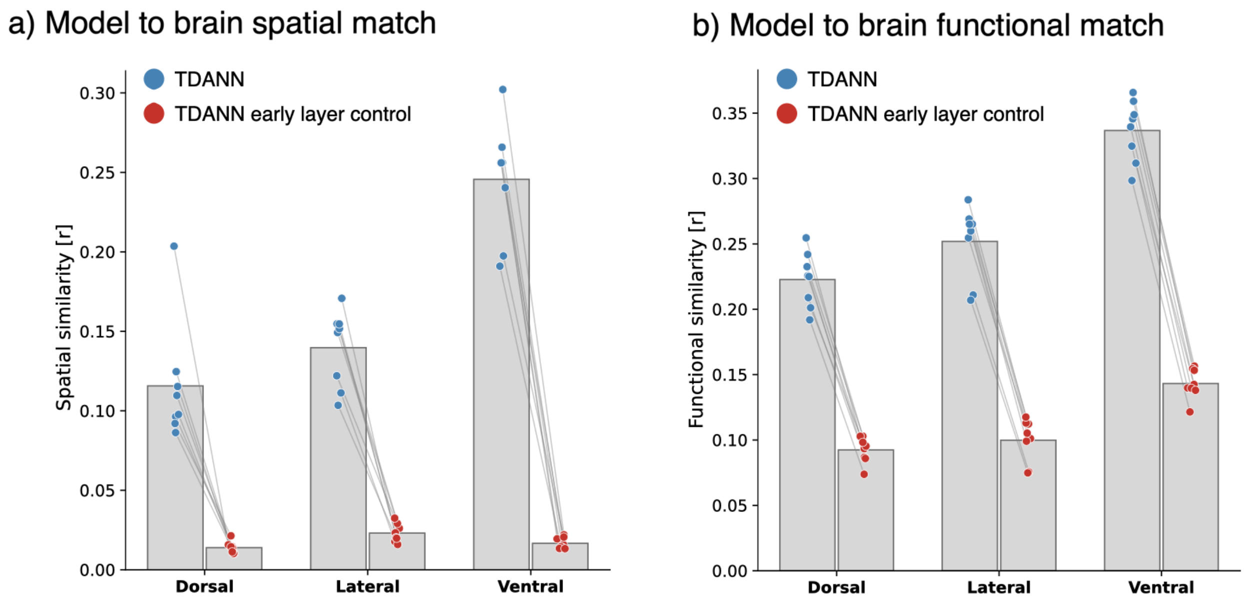


**Figure S5. An earlier TDANN layer does not reproduce the spatial or functional organization of higher-level visual cortex.** We repeated the 1-to-1 model-to-brain mapping using an earlier TDANN layer control (layer 2.0) from the same self-supervised spatial constraints model trained with $\alpha = 0.25$, and compared it to the late TDANN layer used in the main analyses (layer 4.1). **(a)** Model-to-brain spatial match. **(b)** Model-to-brain functional match. Across dorsal, lateral, and ventral stream ROIs, the earlier layer shows markedly weaker spatial and functional correspondence to the brain than the late TDANN layer. Gray bars: mean across subjects; colored points: individual subjects averaged across hemispheres; gray lines connect paired subject values. Thus, the smooth, spatially coherent mappings observed for the late TDANN are not an inevitable consequence of the architecture or the 1-to-1 assignment procedure, but depend on later, higher-level representations emerging in the model. Source data are provided as a Source Data file.

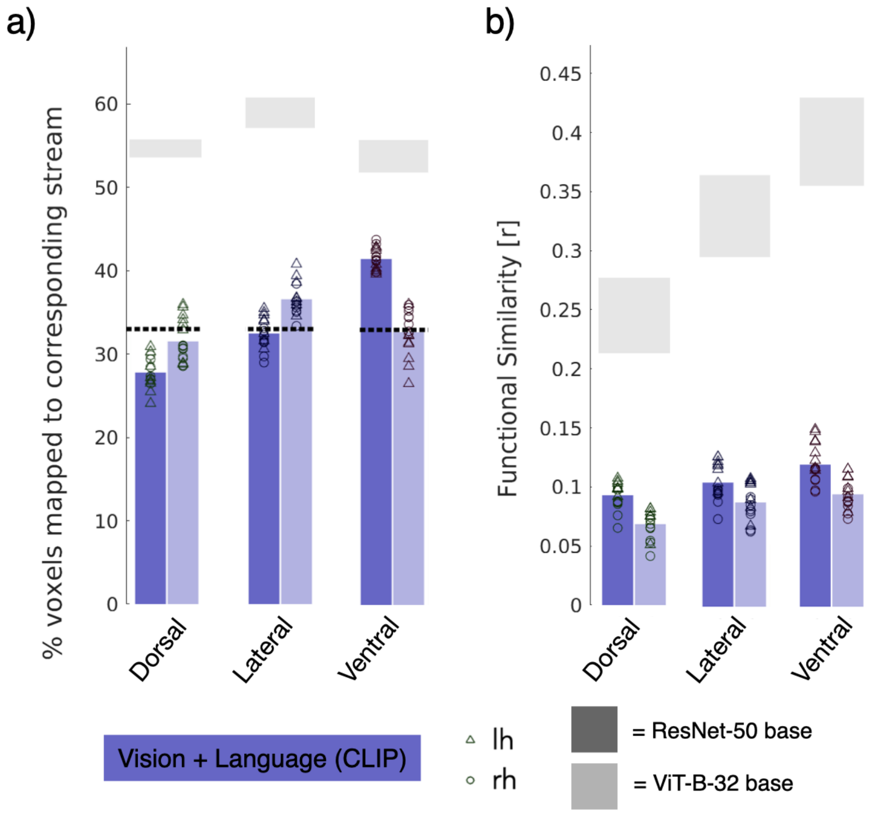


**Figure S6. Minimal differences in correspondence between ResNet-50 and ViT-B-32 versions of CLIP.** We test an additional version of the MB v2 model using CLIP trained with a ViT-B-32 base architecture to compare against the ResNet-50 base version. As in [8], the differences across architectures are minor. However, we do see a slightly increased spatial correspondence to the hypothesized lateral stream, with decreased correspondence to ventral as we move from ResNet-50 to the transformer ViT-B-32 base. While these differences are small, this is interesting in light of our finding that the transformer-inspired, ConvNext-T model in MB v2 shows a significantly lower than chance spatial match to ventral, despite training on object categorization (Fig. 4). Functionally, the ViT-B-32 version provides a slightly poorer functional match overall. **(a)** CLIP model (from multiple behaviors model v2) to brain spatial match: proportion of units assigned to stream by training task. Horizontal dashed line: 33 % chance level. **(b)** CLIP model (from multiple behaviors model v2) to brain functional match: functional similarity of units assigned to stream by training task. For (b) and (c) colored bars: mean across participants and hemispheres; bars are colored lavender for the vision+language training task, dark bars: ResNet-50 base architecture, light bars: ViT-B-32 base architecture; horizontal gray bars: mean noise ceiling across participants and hemispheres ± SD. Source data are provided as a Source Data file.

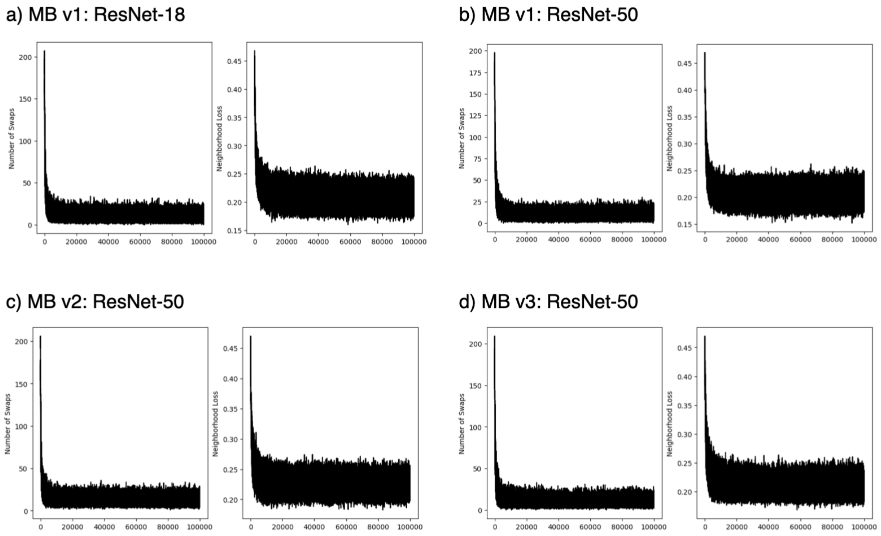


**Figure S7. The neighborhood-preserving swap optimization converges for all multiple behaviors model variants.** To allow the multiple behaviors (MB) models to be compared to the brain using the same simulated cortical sheet framework, model units were assigned positions and then pre-optimized using the neighborhood-preserving swap procedure described in Methods. Here we track the number of swaps accepted (left in each panel) and neighborhood loss (right in each panel) across the 100,000 swap-optimization iterations for **(a)** MB v1: ResNet-18, **(b)** MB v1: ResNet-50, **(c)** MB v2: ResNet-50, and **(d)** MB v3: ResNet-50. Across all model variants, both the number of swaps and the neighborhood loss decrease rapidly and then stabilize, indicating that the procedure reaches a steady solution. Thus, the failure of the multiple behaviors models to recover aligned cortical topographies is unlikely to be explained by lack of convergence of the swapping procedure.

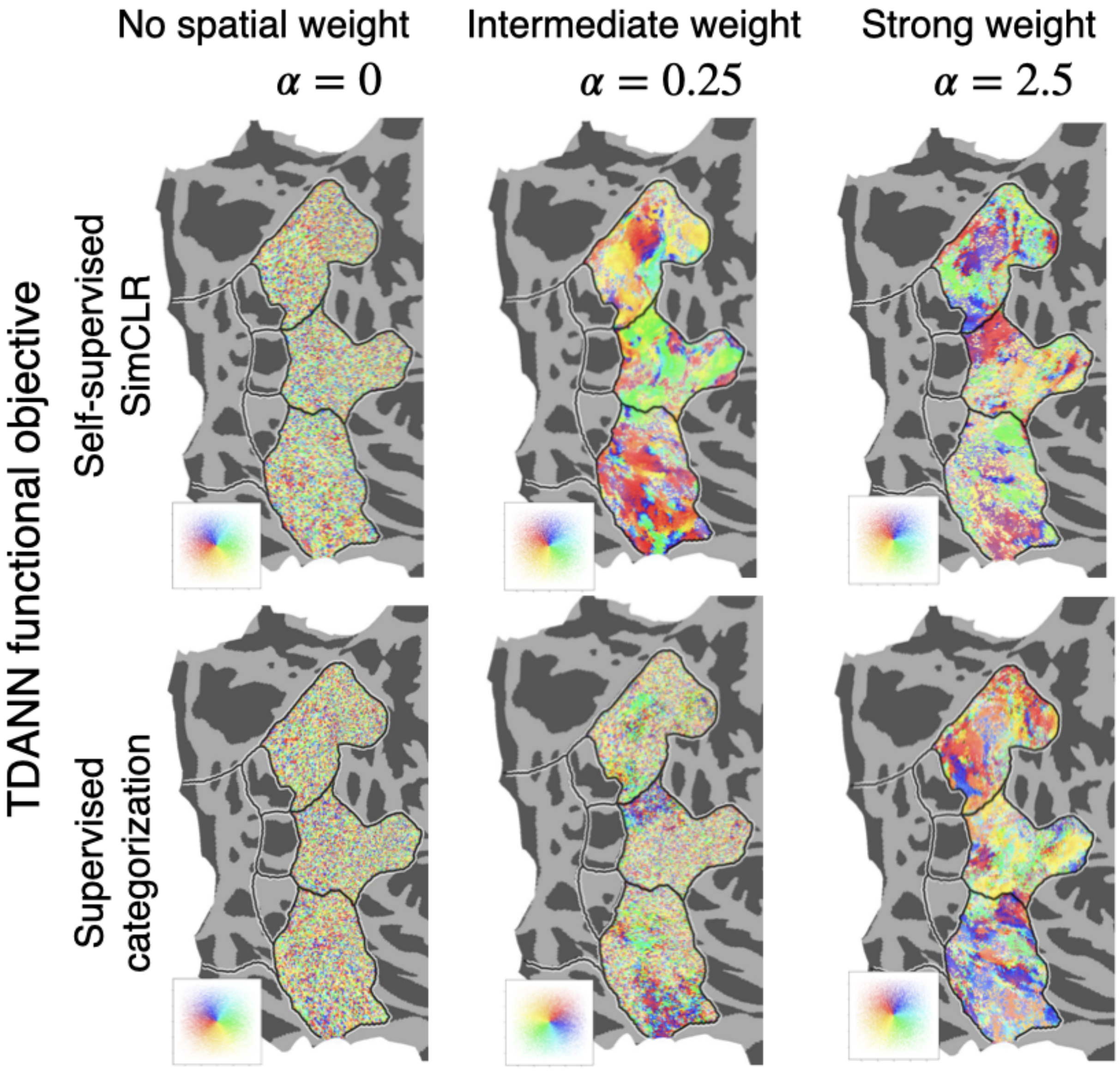


**Figure S8. Mapping between spatial constraints models and the brain for an example flattened right hemisphere using a continuous gradient.** Voxels are colored by location on the simulated cortical sheet (see inset). This rainbow gradient color scheme is "stream-agnostic" in that it does not presuppose the existence of three streams, yet stream clustering emerges. Left: spatial constraints models trained with no spatial weighting ($\alpha = 0$); Middle: spatial constraints models trained with $\alpha = 0.25$, best functional similarity for self-supervised spatial constraints models; Right: spatial constraints models trained with $\alpha = 2.5$, best functional similarity for spatial constraints models trained on categorization.

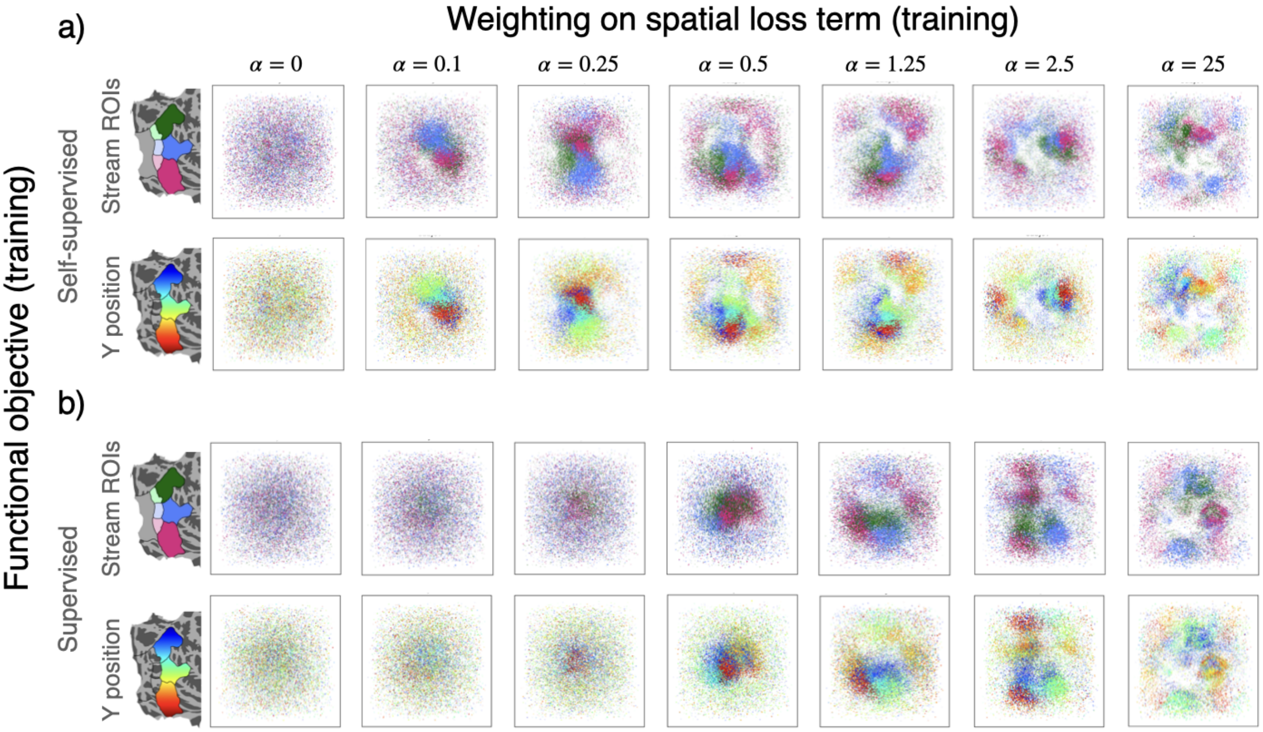


**Figure S9. Model-to-brain 1-to-1 mapping visualized on the simulated cortical sheet of the last convolutional layer of the spatial constraints models.** TDANN model-to-brain mapping visualized on the simulated cortical sheet, for TDANNs trained using the self-supervised, SimCLR task (a) or the supervised, object categorization task (b). This visualization is the reverse of the visualization in Fig. 3a and Fig. 5a of the main text. Here, each square panel shows model units (each dot is a model unit) on the simulated cortical sheet. Units are colored based on the spatial location of their assigned voxel in an example target brain. Opacity of the units reflects the strength of the model-to-brain correlation between responses to NSD images and units are colored by stream (top) or superior-to-inferior spatial gradient (y-position in flat map, bottom). This second color scheme is "stream-agnostic" in that it does not presuppose the existence of three streams, yet stream clustering emerges at self-supervised $0.25 \leq \alpha \leq 0.5$. Each column displays the model-to-brain mapping for an example subject and model seed for a particular spatial weight.

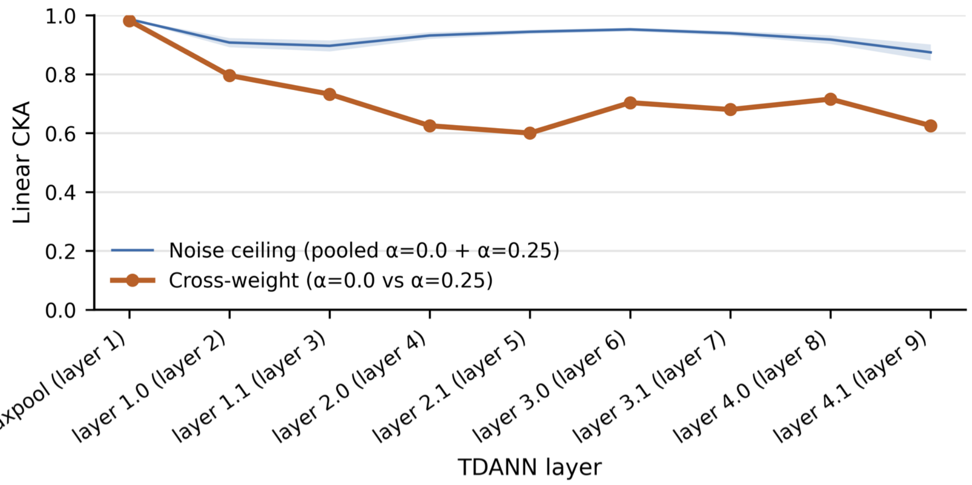


**Figure S10. Adding the spatial loss changes TDANN representations throughout the network.** To test whether the spatial constraint changes TDANN representations, rather than only their spatial layout, beyond variability due to random initialization, we computed linear centered kernel alignment (CKA) between corresponding layers of self-supervised TDANNs trained with no spatial loss ($\alpha = 0.0$) and with the best-performing spatial weight ($\alpha = 0.25$). Activations were extracted for the shared subset of NSD images from layers spanning the network (`base_model.maxpool` through `base_model.avgpool`), flattened across units, and mean-centered across stimuli before computing linear CKA. The blue line shows the pooled within-weight ceiling, computed from seed-pair comparisons within $\alpha = 0.0$ and within $\alpha = 0.25$, while the orange line shows cross-weight similarity between $\alpha = 0.0$ and $\alpha = 0.25$ models. Lines indicate the mean across seed-pair comparisons; shaded region: $\pm$ SEM. Cross-weight similarity is near ceiling at the earliest stage (maxpool) but falls substantially below the within-weight ceiling through much of the network, including at layer 4.1, the late layer used for model-to-brain mapping. These results provide additional evidence that introducing the spatial loss changes internal representational geometry throughout the TDANN, rather than simply rearranging similar units on the simulated cortical sheet. Source data are provided as a Source Data file.

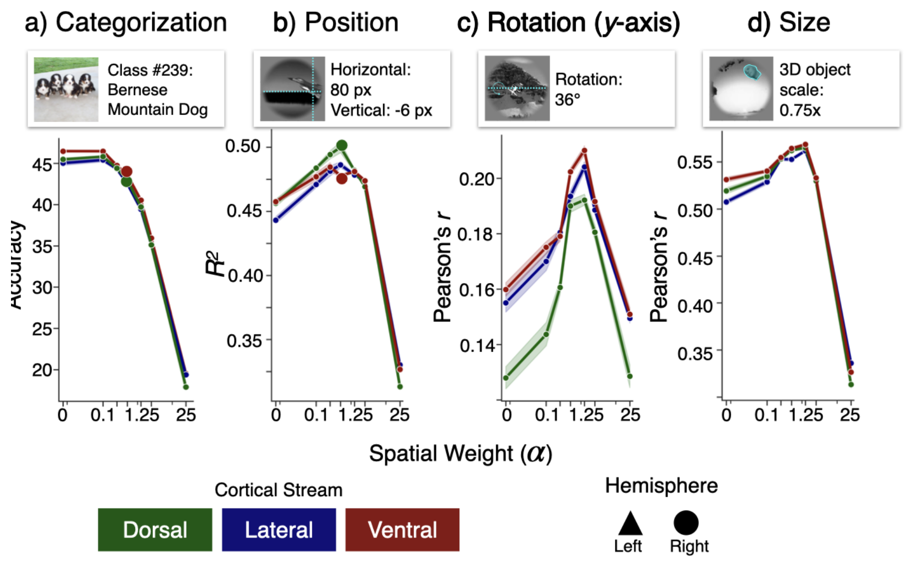


**Figure S11. Adding the spatial loss component to the spatial constraints models during training can improve later transfer performance for some tasks.** Transfer performance for units from the self-supervised spatial constraints models across a range of weightings ($\alpha$) on the spatial loss function for four tasks: object categorization, object position estimation, object pose estimation (y-axis rotation), and object size estimation. For three of the four tasks: position, pose, and size estimation, the addition of a spatial loss term improves performance, with peak performance in the same range of weighting on the spatial loss term that leads to best correspondence with the brain ($0.25 \leq \alpha \leq 1.25$). Units were divided by their assigned stream and then assessed on transfer performance using supervised linear readouts. Results are averaged across model seeds (5), subjects (8) and hemispheres (2) totaling 80 models per point, with the exception of (a) where results represent only one model seed, due to compute constraints. Shaded error bar: ± SE. Larger circles indicate model by stream combinations evaluated main text Figure 6. Source data are provided as a Source Data file.

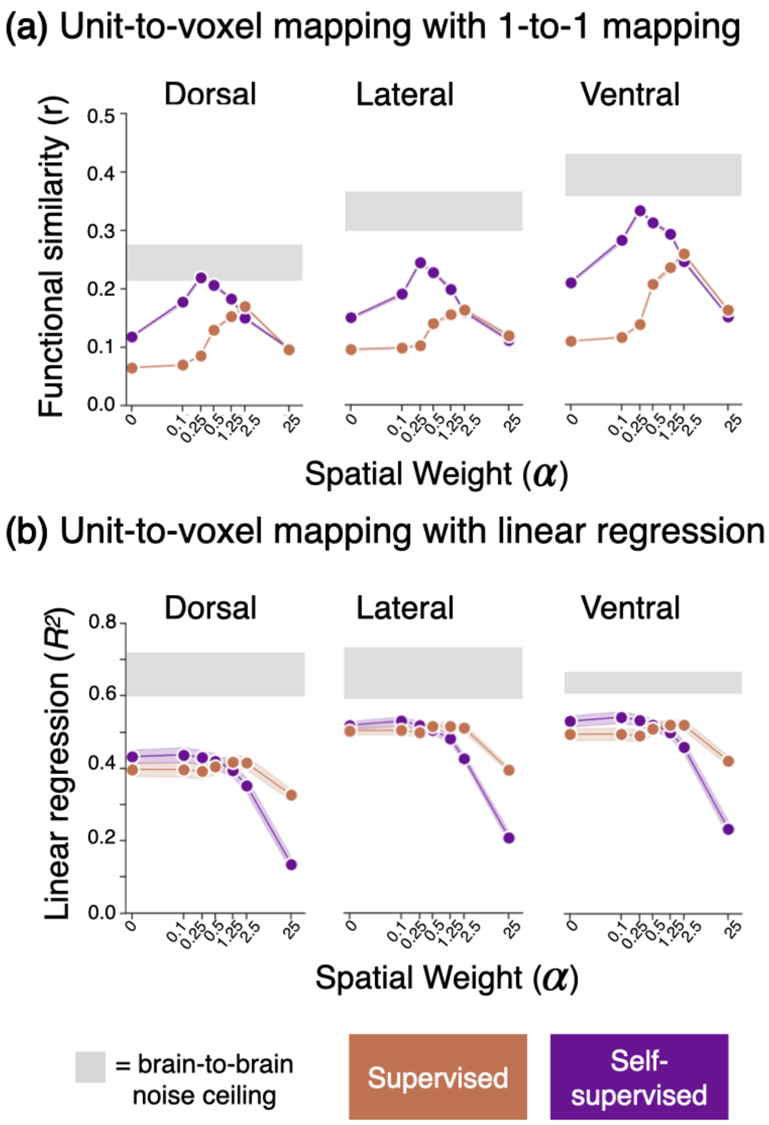


**Figure S12. Mapping between spatial constraints model units and voxels for each stream using either the 1-to-1 mapping (a) or linear (ridge) regression (b).** Functional similarity ($r$) under the 1-to-1 mapping or variance explained ($R^2$) under a linear regression mapping (ridge regression) between model units and voxels for each stream, as a function of the weighting on the spatial constraint and the functional training objective. $R^2$ values are normalized by the noise ceiling of each voxel such that 1.0 corresponds to the intrinsic data noise ceiling. Values are averaged across model seeds and hemispheres; shaded region: SD across participants. Purple: self-supervised spatial constraints model (SimCLR); Orange: supervised spatial constraints model (categorization). Horizontal gray bar: noise ceiling, mean ± SD.

While calculating functional correspondence using linear regression, a commonly-used, less strict mapping of model-units-to-brain, estimates a higher functional correspondence, it critically masks the effects of training task and spatial constraint. In fact, the improved functional correspondence to the brain between TDANNs trained with biologically-plausible self-supervised training and models trained on supervised object categorization nearly vanishes when models are evaluated using linear regression. Of note, these analyses also serve to replicate the canonical finding that when evaluated using linear regression, object categorization trained DANNs are highly predictive of ventral stream neural responses, as the supervised models trained with no spatial weight ($\alpha = 0$) in this analysis are equivalent to off-the-shelf object categorization trained ResNet 18 models. As in previous studies [30, 44], we see noise-corrected variance explained for the ventral stream around 50% for both the supervised object categorization models and self-supervised models trained using SimCLR (mean $\pm$ SD: categorization = $49\% \pm 4\%$, simCLR = $52\% \pm 4\%$). While exact numbers are not reported in [44], from Fig. 2B average Pearson correlations for IT appear to be around roughly 0.75 for both the ResNet-18 models trained using SimCLR and supervised object categorization. This corresponds to variance explained of 56%, a highly similar magnitude to our reported results here using NSD. Source data are provided as a Source Data file.

6. Shaoqing Ren, Kaiming He, Ross Girshick, and Jian Sun. Faster r-cnn: Towards real-time object detection with region proposal networks. *Advances in Neural Information Processing Systems*, 28, 2015.

12. Xizhou Zhu, Weijie Su, Lewei Lu, Bin Li, Xiaogang Wang, and Jifeng Dai. Deformable detr: Deformable transformers for end-to-end object detection. *arXiv preprint arXiv:2010.04159*, 2020.

13. Zigang Geng, Ke Sun, Bin Xiao, Zhaoxiang Zhang, and Jingdong Wang. Bottom-up human pose estimation via disentangled keypoint regression. In *Proceedings of the IEEE/CVF conference on computer vision and pattern recognition*, pages 14676–14686, 2021.

14. Lihe Yang, Bingyi Kang, Zilong Huang, Zhen Zhao, Xiaogang Xu, Jiashi Feng, and Hengshuang Zhao. Depth anything v2. *Advances in Neural Information Processing Systems*, 37:21875–21911, 2024.

15. Kevin S Weiner and Kalanit Grill-Spector. Neural representations of faces and limbs neighbor in human high-level visual cortex: evidence for a new organization principle. *Psychological Research*, 77(1):74–97, 2013.

16. Moritz Wurm and Alfonso Caramazza. Action and object representation in the ventral" what" stream. *PsyArXiv*, 2021.

17. David Pitcher and Leslie G Ungerleider. Evidence for a third visual pathway specialized for social perception. *Trends in Cognitive Sciences*, 2020.

18. Leyla Isik, Kami Koldewyn, David Beeler, and Nancy Kanwisher. Perceiving social interactions in the posterior superior temporal sulcus. *Proceedings of the National Academy of Sciences*, 114(43):E9145–E9152, 2017.

19. Melvyn A Goodale and A David Milner. Separate visual pathways for perception and action. *Trends in Neurosciences*, 15(1):20–25, 1992.

20. A David Milner and Melvyn A Goodale. Visual pathways to perception and action. *Progress in Brain Research*, 95:317–337, 1993.

21. Eshed Margalit, Hyodong Lee, Dawn Finzi, James J DiCarlo, Kalanit Grill-Spector, and Daniel LK Yamins. A unifying framework for functional organization in early and higher ventral visual cortex. *Neuron*, 112(14): 2435–2451, 2024.

22. Jonas Kubilius, Martin Schrimpf, Aran Nayebi, Daniel Bear, Daniel LK Yamins, and James J DiCarlo. Cornet: modeling the neural mechanisms of core object recognition. *bioRxiv*, 2018.

23. Wei Liu, Dragomir Anguelov, Dumitru Erhan, Christian Szegedy, Scott Reed, Cheng-Yang Fu, and Alexander C Berg. Ssd: Single shot multibox detector. In *Computer Vision–ECCV 2016: 14th European Conference, Amsterdam, The Netherlands, October 11–14, 2016, Proceedings, Part I 14*, pages 21–37. Springer, 2016.

24. M. Everingham, L. Van Gool, C. K. I. Williams, J. Winn, and A. Zisserman. The PASCAL Visual Object Classes Challenge 2012 (VOC2012) Results, 2012.

25. Priya Goyal, Quentin Duval, Jeremy Reizenstein, Matthew Leavitt, Min Xu, Benjamin Lefaudeux, Mannat Singh, Vinicius Reis, Mathilde Caron, Piotr Bojanowski, et al. Vissl, 2021.

26. Daniel J Felleman and David C Van Essen. Distributed hierarchical processing in the primate cerebral cortex. *Cerebral Cortex*, 1(1):1–47, 1991.

27. Ilya Loshchilov and Frank Hutter. Sgdr: Stochastic gradient descent with warm restarts. *arXiv preprint arXiv:1608.03983*, 2016.

39. Dawn Finzi, Daniel LK Yamins, Kendrick Kay, and Kalanit Grill-Spector. Do deep convolutional neural networks accurately model representations beyond the ventral stream? In *2022 Conference on Cognitive Computational Neuroscience*, 2022.

48. Yudi Xie, Weichen Huang, Esther Alter, Jeremy Schwartz, Joshua B Tenenbaum, and James J DiCarlo. Vision cnns trained to estimate spatial latents learned similar ventral-stream-aligned representations. *arXiv preprint arXiv:2412.09115*, 2024.
